# Integrated Transcriptomic and Metabolomic Profiling of Sheep Ovarian Tissues Confer their association in Fecundity associated Pathways

**DOI:** 10.1101/2023.12.19.572136

**Authors:** Salsabeel Yousuf, Waqar Afzal Malik, Hui Feng, Tianyi Liu, Lingli Xie, Xiangyang Miao

**Affiliations:** State Key Laboratory of Animal Biotech Breeding, Institute of Animal Sciences, Chinese Academy of Agricultural Sciences, Beijing, 100193, China; Agricultural Genomics Institute at Shenzhen, Chinese Academy of Agricultural Sciences, Shenzhen, 518120, China

**Keywords:** ovarian tissue, metabolomic profiling, positive/negative ion mode, combined ion mode, expression profiling, differentially expressed metabolites, differentially expressed genes

## Abstract

**Background:** Low fertility is considered the major constraint in sheep rearing industry depending on several factors like, estrus cycle, ovulation rate and litter size but fecundity of ewe plays a key role in sheep reproduction, influenced by several intrinsic and extrinsic factors. However, genetic improvements of traits associated with reproduction through conventional breeding is a very complex and slow process. In current study, we went through a comprehensive integration of high throughput transcriptomic and metabolomics approaches to understand the role of key regulatory genes and metabolites in fecundity of two different and widely raised sheep breeds (Small Tail Han & Dolang) in different regions of China.

**Result:** UPLC/MS/MS system based metabolomic profiling of ovarian tissue from both breeds results into the identification of 1,423 metabolites, including 542 DEMs (379 upregulated and 163 downregulated). Integration of metabolomics and transcriptomics data identified 48 pathways contributed by 37 genes and 85 metabolites through regulatory network analysis. Functional enrichment analysis showed significantly enriched pathways associated with fecundity including Riboflavin metabolism, xenobiotics, bile acid biosynthesis, and Drug metabolism, which produces hormones for regulation of ovarian function, ovulation, and establishment of pregnancy. Further, analyzed two restrictive constrained plots analyzed via multivariate statistical analysis. In one plot complement component C3 associated with Leukotriene D4, and Uridine 5’-diphosphate involved in the processes of Neuroactive legend receptor interaction pathway and in second plot IFNGR1 associated with Progesterone, Fumaric acid, and Cortisone involved in the processes of cancer pathway and any disruptions in hormonal balance may induce cancer, which can affect fertility, menstrual cycles, and overall reproductive health.

**Conclusion:** Expression profiling, functional enrichments, co-expression network analysis and integrated transcriptome-metabolome data showed gene-metabolite association in energy metabolism, Inflammation, and drug metabolism, all of which play a role in ovarian physiology and ovarian metabolic disorders. Identification and validation of genes, metabolites, and gene-metabolite interactions will help to elucidate the regulatory mechanisms and pathways underlying sheep fecundity and could be leveraged to improve reproductive traits.

**Graphical Abstract: Scheme of Study:** 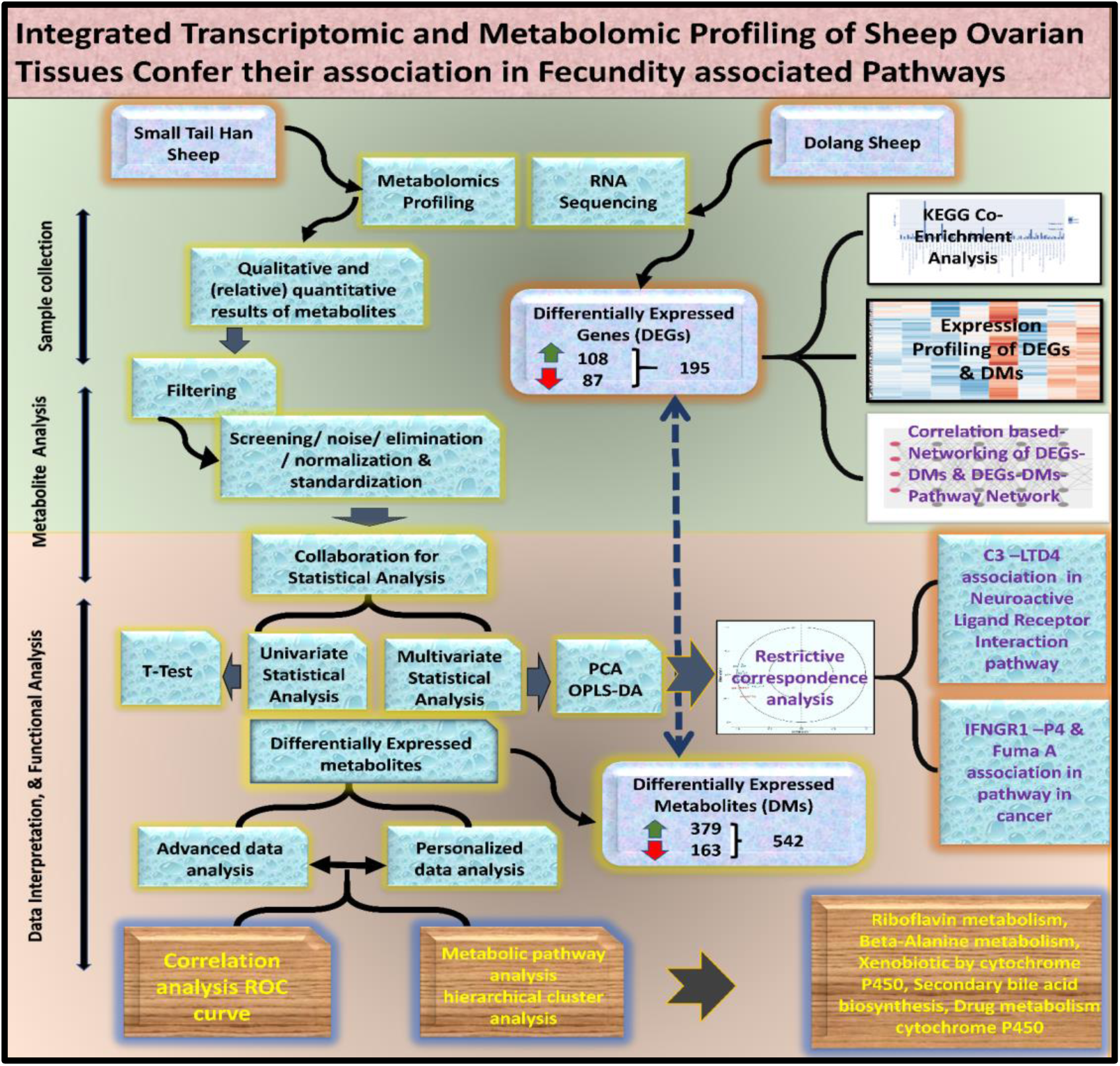

## 1. Introduction

Global food security and sustainability of food production and consumption greatly depends on how to manage livestock production and animal source food consumption. Genetic, nutritional and other environmental factors affect prolificacy traits in sheep. Sheep (*Ovis aries*) with its multi-facet utility for wool, meat, milk, skins and manure is an important livestock and plays a vital role in the agrarian economy [1, 2]. The growing demand for sheep and goat meat products, in recent past, has been driven mainly by animal science and technology, especially regarding production systems, slaughter procedures and carcass fabrication and grading, meat processing, food quality and safety as well consumer preferences and satisfaction [3]. It has been a tough challenge for researchers and breeders to meet the sheep meat requirements so improving the fertility of sheep is an important goal in sheep breeding as it greatly increases the productivity. Modern animal breeding and quantitative genetic studies have shown that traditional selective breeding for particular traits has been slow, with low genetic gains [4, 5]. These limitations of traditional selection methods, which are mainly based on phenotypic characteristics, have led to a growing interest in the identification and characterization of key genes and genomic regions that regulate these economically important traits. The low heritability and high individual variation of fertility traits have been crucial limitations in improving the reproductive performance of sheep breeds [6].

Low fertility, as a major constraint in sheep rearing industry, depends on several factors such as estrus cycle, ovulation rate and litter size, etc. Among them, the fecundity of ewe plays a key role in sheep production, influenced by several factors e.g.; age, heat stress, lambing season, and nutrition states [7, 8]. Ovaries, are the site for folliculogenesis, production of luteinizing hormone (LH) and follicle stimulating hormone (FSH), regulated by non-reproductive organs (central nervous system and pituitary glands)[9–13]. Previously, due to the low heritability of reproductive traits, researchers conducted several studies to improve genetic selection of complex traits, such as higher growth rates, fertility traits, and enhanced disease resistance to obtain responsible candidate genes [14] through genomic studies[11, 15–21]. Although, transcriptomic studies provide insights into molecular regulation of target traits at transcriptional level but the DNA sequence is not the only thing to connect the gene and gene products, however the fingerprint of a product could be characterized by its metabolites content [22]. Metabolites are different types of small biomolecules present in diverse locations widely with varying concentrations [23] that play crucial roles in various cellular and molecular processes determined by specific enzymes and transporters dispersed across tissues and organs [24–26]. Previous studies reported their key role in regulating sheep ovarian physiology and immunity. For example, the levels of certain hormones, such as estrogen and progesterone, are regulated by the metabolic processes in the ovaries [27]. Identification and characterization of biomolecules is therefore challenging, yet also critically important for understanding biology at a molecular level and preventing disease. Metabolomics provides data-rich information on metabolic changes that reflect genetic, epigenetic, and environmental variables influencing cellular physiology [28, 29]. Generally, metabolic profiling provides the small number of metabolites relative to the estimated numbers of genes, transcripts and proteins, however they are of great significance in translational studies. The interpretation of metabolic pathways provides information for biological mechanisms of cell/tissue [30].

Metabolomic studies systematically assess the unique chemical elements produced during specific cellular processes by describing the metabolites profiles of individuals that undergo similar biological processes and transformations [31]. However, RNA-Sequencing (RNA-Seq) analysis interrogates the level of mRNA from an organism to estimate its gene expression profile at a given moment [32]. Identification of active and repressed genes in different regimens would lead to the discovery of the biological pathways affected by these functional differences. In this study, Ultra Performance Liquid Chromatography and tandem mass spectrometry (UPLC-MS/MS) system based metabolomics approach was used for the first time to analyze the ovarian metabolites from two most famous sheep breeds with opposing characteristics (Small-tailed Han and Dolang sheep) native in two different regions of China (Shandong Province and south Xinjiang) respectively with fecundity variations and performed transcriptome analysis on the same corresponding group to examine and elucidate various complex physiological and biological processes related to their fertility [33–35]. Small Tail Han sheep (X) is a medium wool, fairly small with ewes weighing 35–45 kg on average, is known for its very high rates of reproduction and extremely high fecundity (about 229%) made it a growing part of China’s livestock sector [36]. They average 3.44 lambs a year per ewe, with an average litter size of 2.29 [12] and under ideal conditions, each ewe can produce 9 lambs every 2 years [11, 37, 38]. Though, Dolang sheep (D) are widely bred in the south Xinjiang region of China, high litter rate and strong adaptability but low fecundity compared to Small Tail Han sheep [11, 39]. In the past few years, molecular regulatory mechanisms for higher prolificacy, year-round estrus, and adaptability in health and disease in X and D sheep have been studied through high throughput sequencing technology [11, 40, 41], the lack of metabolic profiling and integration with transcriptomics from ovine tissue of Dolang sheep and Small Tail Han sheep, limits further understanding. Thus, aimed to analyze the fecundity of sheep by multi-omics approaches. In present study, we performed metabolomics profiling of ovarian tissue, employing UPLC/MS/MS technology to explore differentially expressed metabolites and significantly enriched metabolic pathways that may regulate ovarian function. As well as elucidated the integration of differentially expressed metabolites and differentially expressed genes (resulting of transcriptomics), to identify gene-metabolites association influencing sheep fecundity. Hence, integration of metabolomics and transcriptomics generates hypotheses about metabolite–gene interactions might have functional relevance in signaling pathway that regulate ovarian function. Our methodology generates physiologically meaningful results will contribute in understanding of the underlying mechanisms of sheep fertility.

## 2. Materials and Methods

### 2.1. Experimental animals

Experimental animals of this study were approved by the head of Animal Care at the Institute of Animal Science, Chinese Academy of Agricultural Sciences. The experiment was performed in accordance with the guidelines and regulations for the care and use of laboratory animals established by the Ministry of Agriculture of the People’s Republic of China. Samples were collected from Small Tail Han Sheep and Dolang Sheep ovary tissue (n=6 for each breed), and immediately kept in liquid nitrogen (−80°) for long-term preservation and further extraction at Chaoyang district, Beijing, China.

### 2.2. Metabolomics profile

12 ovarian samples (D (n = 6), X (n = 6)) were placed in lyophilizer (Scientz-100F) for chemical extraction (low molecular weight water-soluble). Then, samples were vacuum freeze-dried, and ground into powder (30 Hz, 1.5 min) using a grinder (MM 400, Retsch). 100 mg of aliquot was taken from each sample and dissolved 1.2 mL of 70% methanol, and then vortexed six times for 30s after each 30 min. Further, we added ceramic beads to ground the samples at 45Hz for 10min, sonicated for 10min (ice water bath) and refrigerated overnight at 4 °C. The samples were centrifuged at 12,000 rpm for 10 min. We carefully collected 500 μL of the resulting supernatant in to EP tube, and aspirated the supernatant. Then, samples were filtered with 0.22μm pore size microporous membrane (0.22 μm pore size) and stored in the injection flask. In addition, samples were sonicated in ice water bath for 10 min at 4 °C and centrifuged (12000 rpm) for 15 min. The resulting supernatant (120 μL) were collected into a 2ml injection bottle for LC-MS analysis. Samples were analyzed after separation through ultra-performance liquid chromatography (UPLC) (SHIMADZU Nexera X2, (https://www.shimadzu.com.cn/) system composed of Waters Acquity I-Class instrument equipped with UPLC HSS T3 column (Agilent SB-C18, 1.8μm, 2.1 mm × 100 mm). For positive and negative ion mode (ESI-MS), the gradient system consisted of mobile phase A (ultrapure water with 0.1% formic acid) and mobile phase B (acetonitrile with 0.1% formic acid), with optimized condition, for example, elution gradient was 5% of phase ratio for 0.00min, capillary voltages for positive and negative ion mode were 2000 V and 1500 V, ion source temperature was 150 °C, cone voltage was 30V, desolvation gas temperature was 500 °C, desolvation gas flow rate was 800L/h, and backflush gas flow rate was 50 L/h. We collected primary and secondary data in MSe mode through Waters Xevo G2-XS QTOF high-resolution mass spectrometer with the MassLynx V4.2. Data acquisition was performed simultaneously in dual channel (low Collison energy and high collision energy). The high Collison energy was 10-40 V and the low collision energy was 2 V, but the scanning frequency of the mass spectrum was 0.2 s.

### 2.3. Data processing of Metabolites

The raw data (MassLynx V4.2) was further processed by Progenesis QI software for peak extraction, peak alignment and other data processing operations. Based on METLIN database of Progenesis QI and self-built database of Baimaike, we determined the theoretical fragmentation and mass deviation within 5ppm. In addition, we performed a series of quality control methods to obtain high-quality data for subsequent analysis. After normalizing the raw peak area with the total peak area, the resulting 3D data were imported into the R-3.1.1 using the following Packages, ggplot2 and scatterplot3d for Principal component analysis (PCA). Then, we calculated Spearman correlations using R-3.1.1. Package of heatmap was used as an assessment index for samples repeatability within group, between the group and quality control samples. Then, identified compound were annotated with KEGG, Human Metabolome Database (HMDB), and lipid maps databases for pathway information and classification. Based on the grouping information, we calculated and compared the difference multiples, for example, T-test was used to calculate the significance p-values of each compound. The R package (3.32) ropls was used to perform orthogonal partial least square discriminant analysis (OPLS-DA) modeling, and seven-fold cross validation with 200 times permutation tests was used to verify the robustness and reliability, and further validated the model. The variable importance in projection value (VIP) of the model was calculated using multiple cross-validation. To compare the difference multiples in the quantitative information of the metabolites in each group, we used a logarithmic (LOG) converter to process the data by generalized logarithmic transformation (LMGene package in R language). The results were processed by combining the difference multiple, the P value of the t-test, VIP value of the OPLS-DA model and fold change was taken to sort out the differentially expressed metabolites (DEMs) with the strict cutoff of, FC>1, P value<0.01 and VIP >1. Then, we applied multiple variance analysis on differentially expressed metabolites using R tool with correlation (R-3.6.1; Package: corrplot.), Z-score plot (R-3.6.1; Package: ggplot2 scale), k-means clustering (R-3.6.1; Packages: NbClust, ggplot2), volcano plot (R-3.6.1; Package: ggplot2), and cluster analysis (R-3.6.1; Packages: heatmap, RColorBrewer).

### 2.4. Functional Annotation of Differentially expressed Metabolites

We determined metabolic pathway of DEMs by combining R software and KEGG (Kyoto Encyclopedia of Genes and Genomes). In enrichment analysis, the p-value of the differentially expressed metabolites were calculated using a hypergeometric distribution test with R-3.6.1; Package: cluster Profiler [56], enrich plot, and ggplot2.

### 2.5. ROC curve analysis

We performed receiver operating characteristic curve (ROC) analysis to investigate biomarkers in ovarian function [42]. The area under curve (AUC) is a very useful measure of the ROC curve [43]. We separately performed ROC analysis on the screened differentially expressed metabolites in each group, and the AUC results were calculated using the R-3.6.1; Package: MetaboAnalystR [42], caTools, pROC.

### 2.6. Data processing of Transcriptome

We converted the raw image data files obtained by RNA sequencing into raw sequencing sequences through Base Calling analysis and stored them in FASTQ format, including the sequence information of reads and their corresponding sequencing quality information. The raw data quality was determined through Trimmomatic software. Then, we used hisat2 software mapped clean reads with reference genome to find out their location on reference genome and to acquire sequence information specific to the sequencing samples. Then, FPKM values for each gene in each sample were calculated using Feature Counts software v 1.5.0-p3, which is used for the calculation of protein-coding gene expression. The differential gene expression between samples was analyzed by comparing RNA-seq data via DESeq2 (R package v 1.16.1), differences were considered statistically significant at p<0.05 with a fold change > 2.

### 2.7. Metabolomics and the Transcriptomic Integrative Analysis

Finally, DEMs resulting of metabolomics were combined with the DEGs obtained via transcriptome. KEGG annotation was performed for differentially expressed metabolites and differentially expressed genes of the same group. However, we only selected gene and metabolite pathways with p-value < 0.05 for further analysis. The Pearson correlation between genes and metabolites were calculated from the z-value transformation of preprocessed data. The correlation coefficient (CC) and correlation p-value were filtered with the screening threshold: | CC|> 0.80 and CCP < 0.05 to screen out significant differentially expressed genes and differential metabolite association. In addition, the differentially expressed metabolites and differentially expressed genes were classified by K-mean for each group. Then, we achieved a consistent trend between metabolites and genes due to certain relationships and applied heatmap clustering analysis by classifying them. Additionally, we constructed correlation-based network for DEGs and DEMs that screened by the pathway for mapping. Multivariate analysis including canonical correlation analysis (CCA) and typical correlation analysis were performed based on correlation between comprehensive pairs of variables to reflect the overall correlation between two sets of indicators and draw the correlation map.

## 3. Results

### 3.1. LC/MS System Based Metabolomic Analysis and Identification

To provide an overview of the metabolites in ovarian tissue, we performed non-targeted metabolic profiling of sheep ovarian tissue using UPLC/MS/MS system. To check the reproducibility of the analytical procedure, we went through a strict quality control (QC) measure. The repeatability of metabolite extraction and detection was mediated by overlap analysis of the positive and negative ion current in the different QC samples. The results showed the peak retention time and peak areas of the QC samples overlapped in the positive and negative ion current profiles during metabolite detection (**Figure 1A & 1G**). In addition, the positive and negative ions in response of internal standard L-2-chlorophenylalanine in different samples determined the consistent in retention times and peak areas shown in total ion chromatography (TIC) diagram of each sample (**Figure 1B-C-D-E-F & H-I-J-K-L**). This indicates that signal stabilizes at the moment when the identical sample was detected at a different time

**Figure 1.**
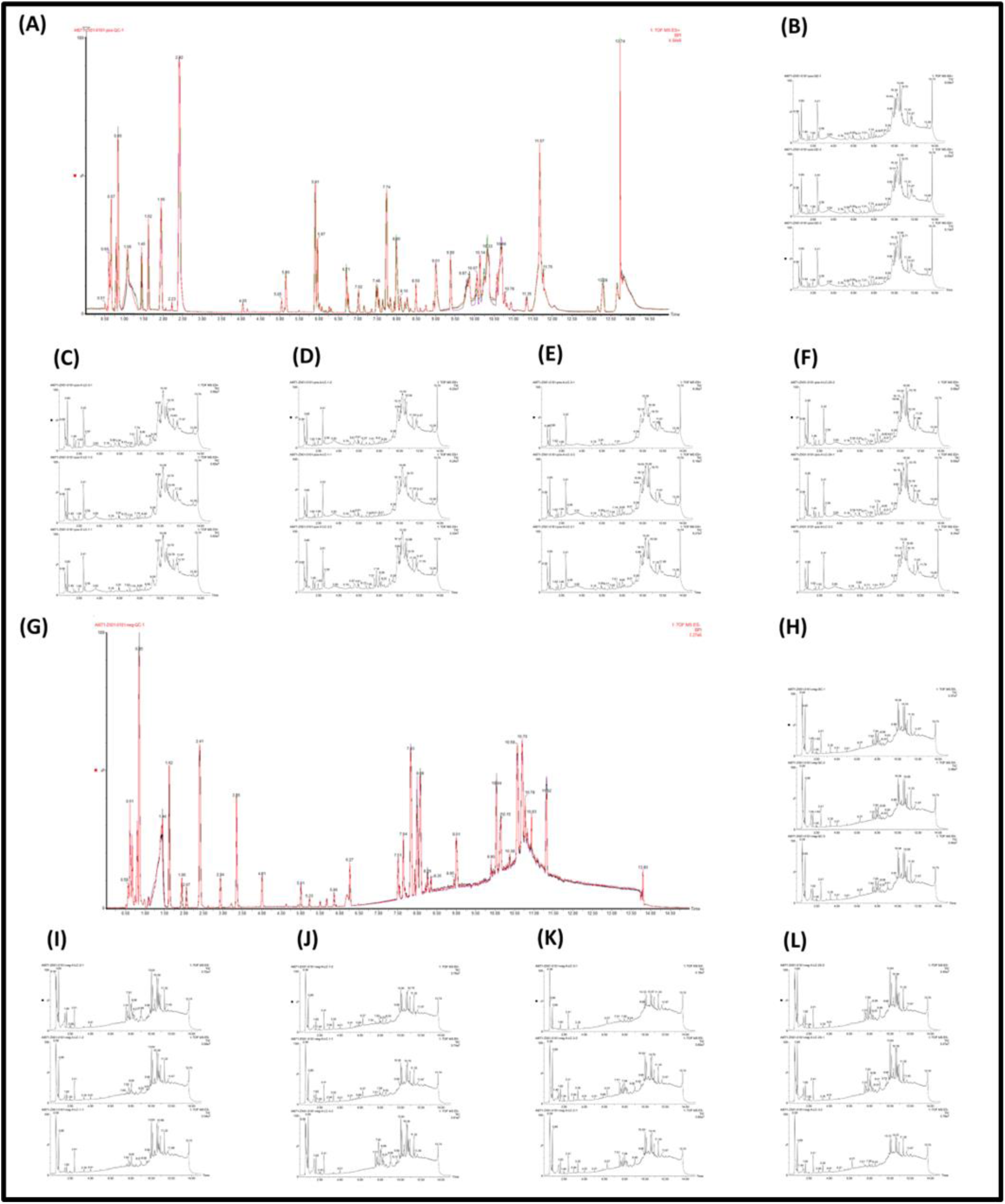
Samples quality control process, instruments stability and TIC plot. (A) Positive ion BPI plot for all QC samples. (B-F) Positive ion mode internal standard response for each sample TIC plot. (G) Negative ion BPI plots for all QC samples. (H-L) Negative ion mode internal standard response for all QC samples.

A total of **1423 metabolites** were identified, **801** metabolites from positive ion mode and **622** from negative ion mode. Instrument stability significantly ensures the repeatability and reliability of our metabolomics data. PCA analysis of samples including QC samples ensured data quality and it was also evident that all data were relatively centralized. The contribution rates of first three primary components (PC1, PC2, PC3) for each sample were 98.8%. The results provide an understanding of overall metabolic differences between sample groups as well as the magnitude of variability between samples within the group, indicated an overlap between biological replicates. The distribution differences within the group were not significant. However, the difference between the groups were significant in negative and positive ion mode (**Figure 2A-C**). This indicates that the data quality was very high. Furthermore, applied PCA on combined (+, −) ion metabolites showed the 95% ellipse confidence interval between biological replicates **(Figure 2E)**. Four biological duplications in X group were concentrated on the right side of the PCA. By contrast, biological duplications in D group were concentrated on the right side of the PCA. In addition, we assessed the correlation among biological replicates using **Spearman Rank Correlation** separately for positive ion mode, negative ion mode, and combined ion (+, −) metabolites. Each of them showed a strong correlation between samples replications (**Figure 2B-D-F**). The correlation coefficients between the X and D groups also showed that X and D has some biological duplications.

**Figure 2.**
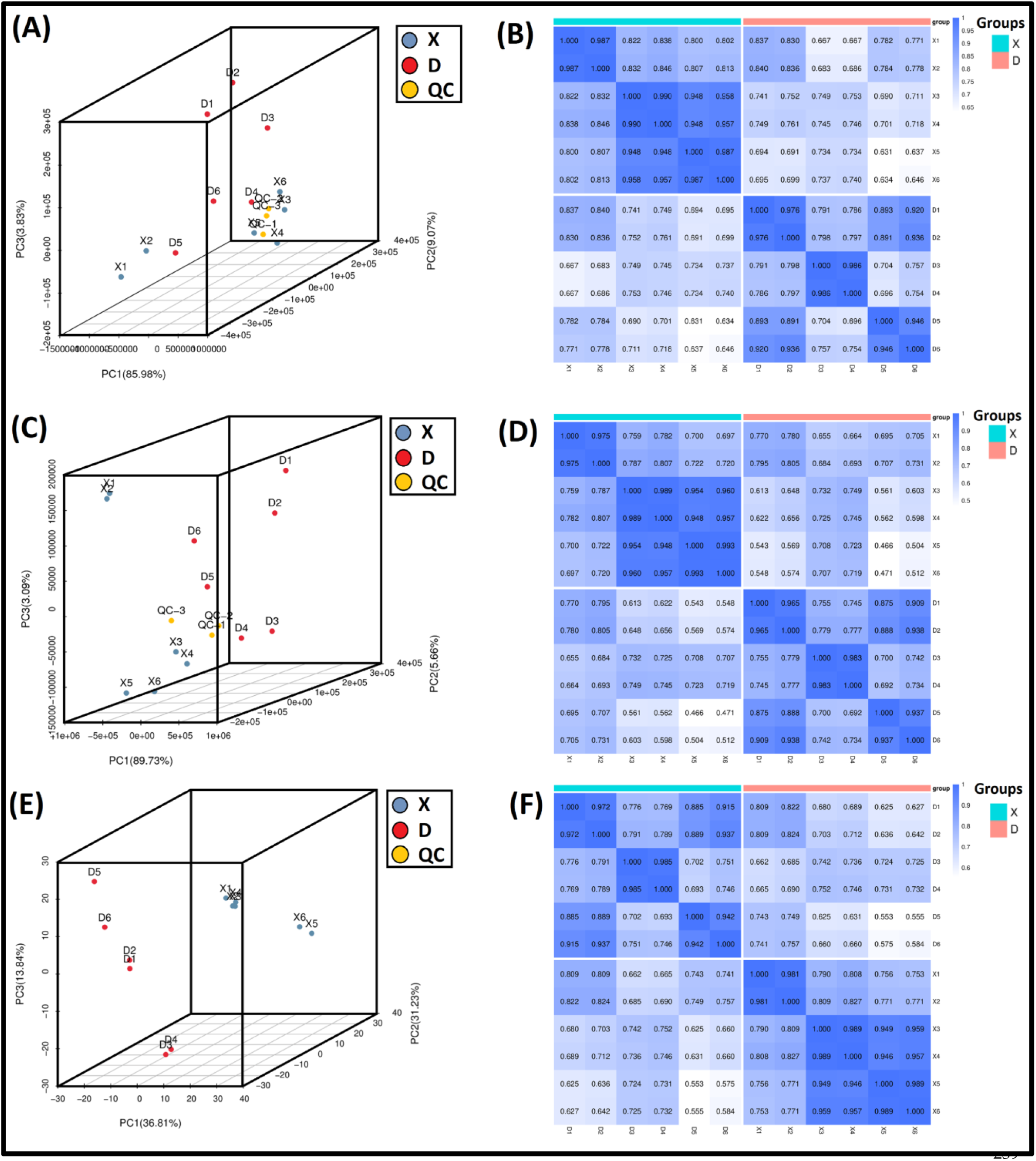
Principal components Analysis and expression of quantitative indicators. (A-B). PCA score plot and spearman correlation plot for Positive ion metabolites. (C-D). PCA score plot and spearman correlation plot for negative ion metabolites in D_vs-X. (E-F) Combined (+, −) ion metabolites PCA and spearman correlation plot.

Only 664 of the 1,432 detected metabolites were annotated with the 254 KEGG pathways. Largest number of metabolites ((n=61) were enriched in the biosynthesis of plant secondary metabolites pathway, followed by ABC transporters (n=48), purine metabolism (n=46), pyrimidine metabolism (n=32) and so on, as shown in **Figure 3A**. Similarly, 842, 368 metabolites were successfully annotated with HMDB and LIPID MAPS. The HMDB database provide detail information about small molecule metabolites found in human body, which are critical for clinical chemistry and for biological biomarker. We determined eleven super classes of HMDB along with HMDB taxonomy as shown in **Figure 3B**. The largest group of metabolites (n=135) were found in the organic acid and derivatives super class, which include carboxylic acid and derivatives hydroxy acids and derivatives taxonomy. LIPID MAPS database classified biolipids into seven broad categories along with several major classes and subclasses. Among seven categories, fatty acyls were leading with maximum metabolites (n=121), followed by glycerophospholipids (n=108), sterol lipids (n=51), and so on as shown in **Figure 3C**.

**Figure 3.**
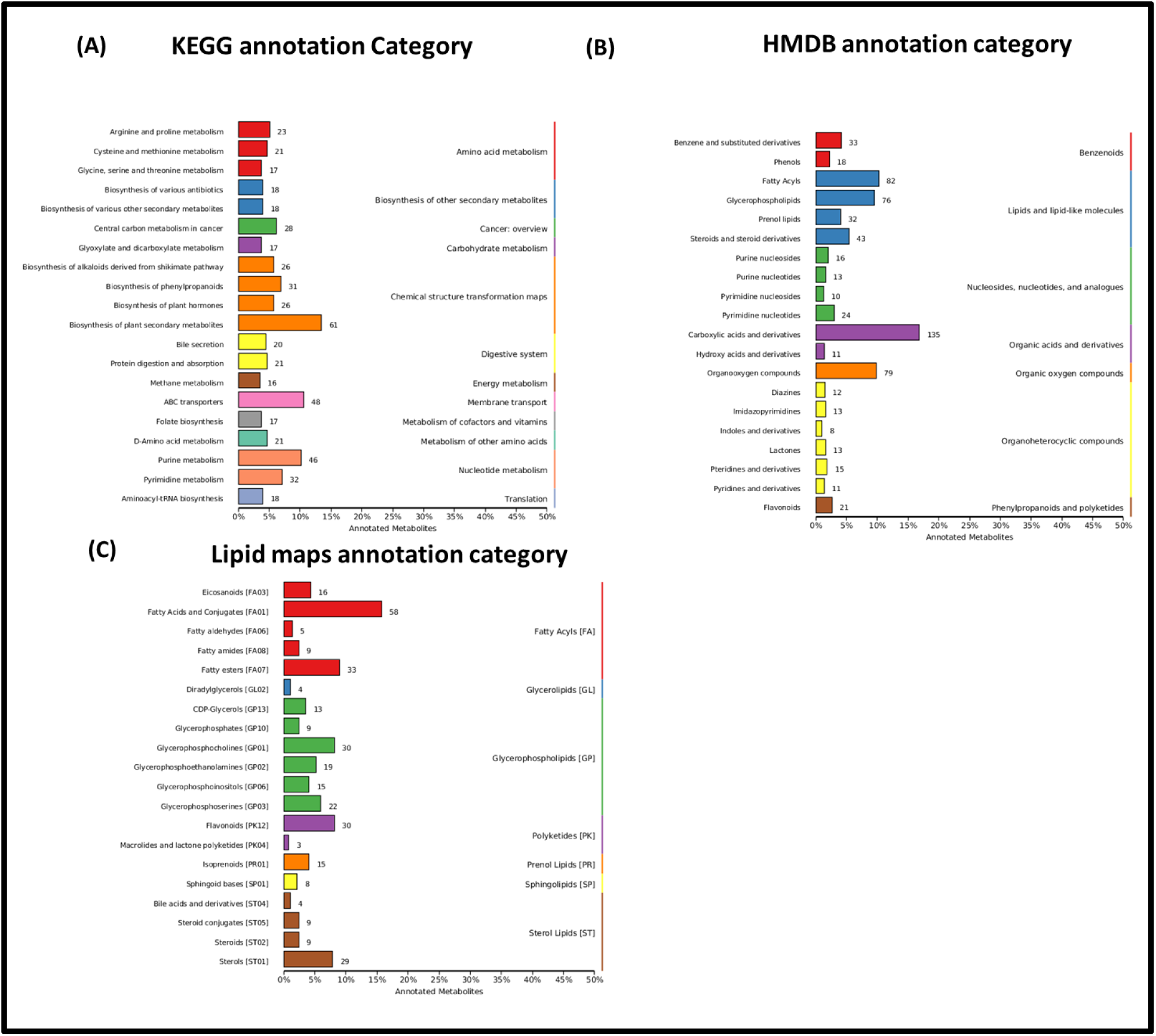
Overall Identified metabolites annotation with (A) KEGG (B) HMDB (C) Lipid Map databases.

### 3.2. Identification of Differentially expressed metabolites (DEMs) and their Pathway Analysis

Before performing variance analysis on the resulting data of combined (+, −) ion metabolites, analyzed PCA plot of differential metabolites obtained at positive ion mode, negative ion mode and combined (+, −) ion mode determined high degree of variability between and within the difference grouping X and D (Figure 4A-B-C). We assessed metabolite differences between groups through the supervised orthogonal projection to latent structures-discriminant analysis (OPLS-DA) model. In OPLS-DA model, score plots for positive, negative and combined (+, −) ion were R2Y = 0.995, and Q2Y=0,963, R2Y = 0.997 and Q2Y=0,966, R2Y = 0.997 and Q2Y = 0.978, as shown in **Figure 4D-E-F**. This indicates that the model was reliable, stable and predictable. A premutation verification confirmed the reliability of OPLS-DA model. The intercepts between the regression line of R2Y and Q2 and the longitudinal axis of the positive, negative and combined (+, −) ion were <0. The fraction of Y variables was gradually increased with the reduction of replacement retention reduced, while Q2 gradually decreased. This indicates that the model is robust without overfitting. All samples were within the 95% confidence interval. These results verified that the D_vs_X was represented different metabolic patterns. Using extended logarithmic transformation, the results of our qualitative and quantitative analyses of the metabolites were compared (R language LMGene package). We aggregated the difference-multiple across biological replicates and selected 542 differentially expressed metabolites (DEMs) using the cutoffs of fold change (FC) > 1, P value 0.01, and VIP >1. The number of metabolites from positive and negative ion mode were 264 and 278 respectively. Based on the expression, 163 down-regulated and 379 up-regulated metabolites were identified in D_vs_X (Figure 5A). The differences in the expression level of these DEMs represented through Volcano Plot showed statistically significant differences in X and D (Figure 5B). Each point in the volcano chart represents a metabolite, the size of the scatter point represents the VIP value of the OPLS-DA model. In Figure 5B, blue dots represent the down-regulated, red dots represent up-regulated DEMs, and the gray dots represent insignificant DEMs (**Supplementary Table S1**).

**Figure 4.**
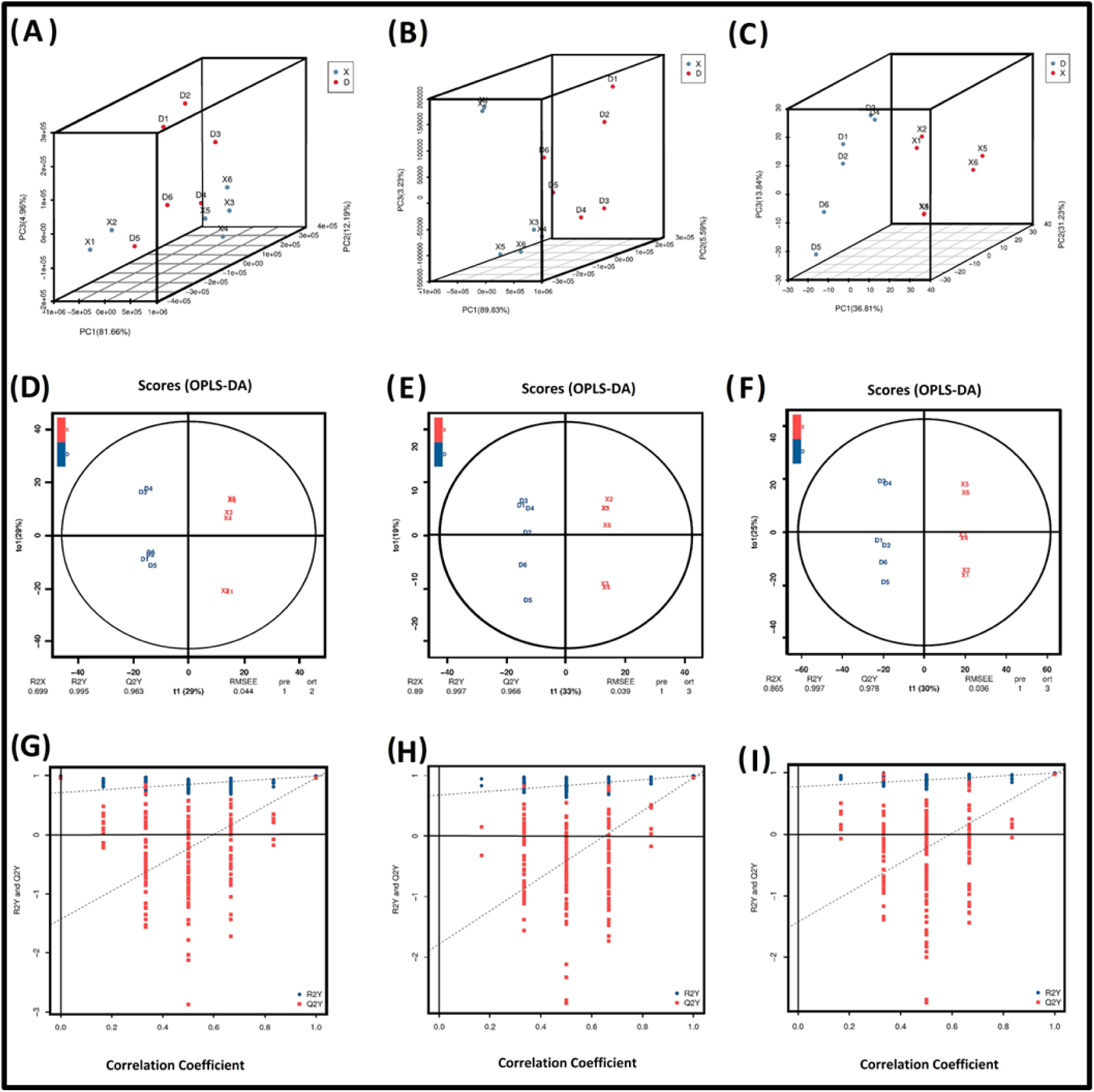
PCA analysis and OPLS-DA SCORE PLOT between samples. (A-C) Positive ion, negative ion, and combined ion (+, −) PCA in D_vs_X. (D-F) OPLS-DA score plot for Positive ion, negative ion, and combined ion (+, −). (G-I) OPLS-DA model verification diagram for Positive ion, negative ion, and combined ion (+, −). The horizontal axis in the figure represents the similarity with the original model, the vertical axis represents the value of R2Y or Q2 (where R2Y and Q2 of 1 are the values of the original model in the abscissa. Note. Both R2Y and Q2 were smaller than the R2Y and Q2 of the original model. All the points on the left side of the figure (displacement test) were lower than the points at the abscissa (the original model) which indicated reliability of the model.

**Figure 5.**
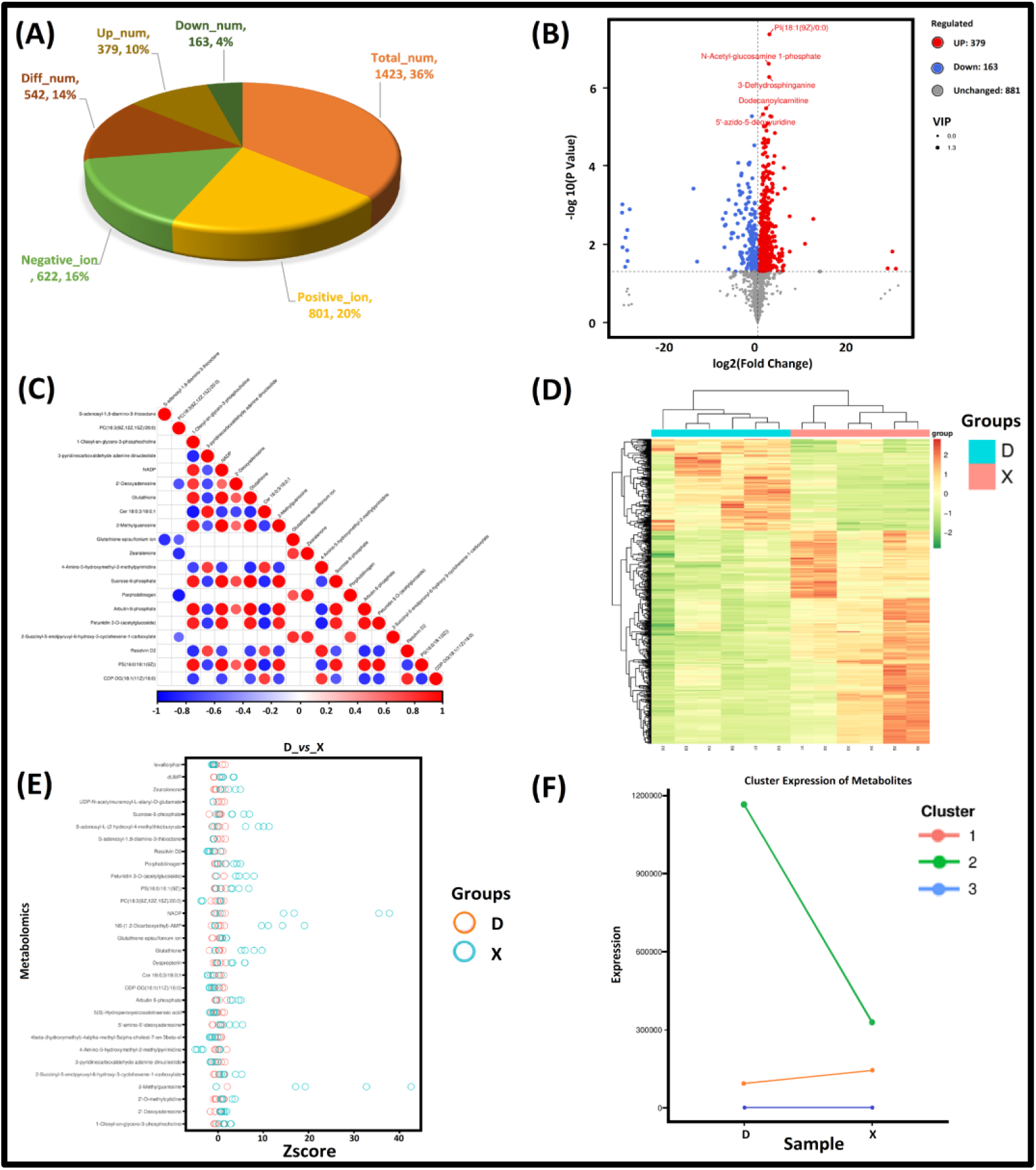
Differentially expressed metabolites from ovary tissue of Dolang sheep (D) and Small Tail Han sheep (X). (A) Overall statistics of metabolites in D_vs_X. (B) Volcano plot of DEMs. (C) Correlation plot based on p-value of top 20 DEMs. (D) Cluster Heat map representation 20 of DEMs expression. (E) Z-score plot representation of DEMs. (F) Differential metabolic union k-means clustering trend chart.

At the same time, from correlation analysis examined the significant synergistic relationship between DEMs with the criteria of p-value ≤0.05 in D_vs_X as shown in Figure 5C & Supplementary Table S2. Further, visualized the expression of DEMs through Heat map cluster analysis **(Figure 5D**). Moreover, Z-score value from quantitative values of metabolites measured the deviation of the experimental group (X) from the control group (D) as shown in **Figure 5E (Supplementary Table 3**). Differential metabolic union k-means clustering used to divide metabolites into three cluster with similar trends based on the nearest mean (Figure 5F), indicating the existence of regulatory relationships within the metabolites of each cluster. The number of DEMs represented by each cluster were 14,1, and 527, respectively.

Additionally, identified DEMs were classified into 49 categories, large number of metabolites (51) fell in the class of carboxylic acids and derivatives, followed by fatty acyl groups (29), organooxygen compounds (29), Glycerophospholipids (27), Steroids and steroid derivatives (23), Benzene and substituted derivatives (14), Pyrimidine nucleotides (14), Flavonoids (10), Pteridines and derivatives (10), Imidazopyrimidines (8), and so on as shown in Supplementary Figure S1.The results of KEGG mapping of DEMs were classified in accordance with the type of pathway in KEGG. Based on rich factor and differential abundance score determined significant pathways which were consistent with the pathways obtained via KEGG enrichment analysis. We observed that majority of the enriched pathways focused on metabolism. By contrast, only a few focused on genetic and environmental information processing. The major enrichment of KEGG pathways included riboflavin metabolism, metabolism of xenobiotics by cytochrome p450, beta-alanine metabolism, secondary bile acid biosynthesis, drug metabolism-cytochrome p450 (**Figure 6A-B-C**), may play potential role in reproduction. Previously, several studies have been reported the contribution of Riboflavin metabolism[44–48] and metabolism of xenobiotics by cytochrome P450[49, 50] in fecundity of sheep. We found several metabolites including (-)-Riboflavin (pos_4, neg_4), Guanosine triphosphate (pos_521), and FAD (neg_170) were mainly enriched in riboflavin metabolism **(Figure 6D).** Members of glutathione family such as; S-(Formylmethyl) glutathione (neg_134), Glutathione episulfonium ion (neg_86), (1R)-Glutathionyl-(2R)-hydroxy-1,2-dihydronaphthalene (pos_411), and S-(Formylmethyl)glutathione (pos_86) were enriched in the metabolism of xenobiotics by cytochrome P450 pathway.

**Figure 6.**
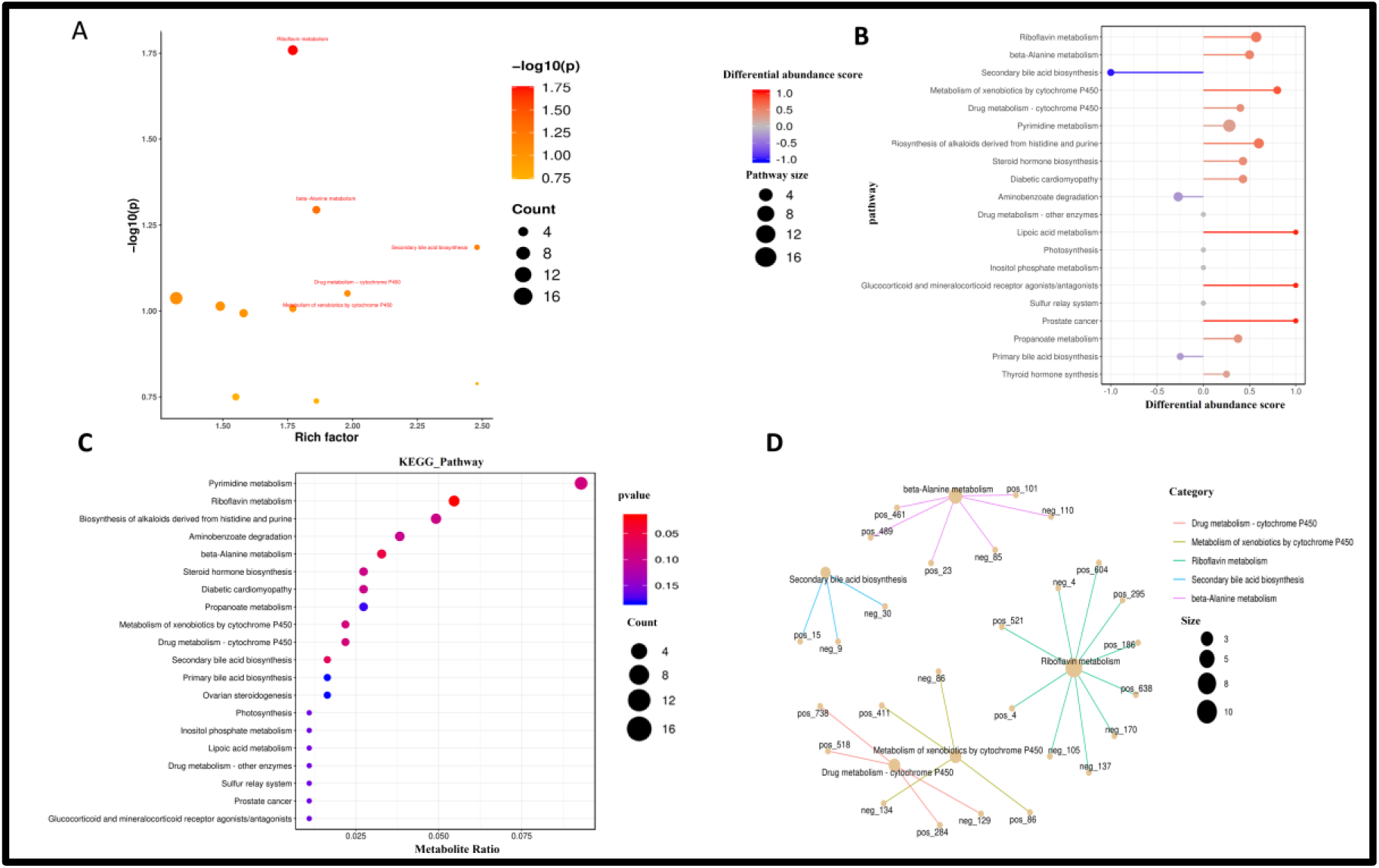
(A-C) Differentially expressed metabolites functional annotation, KEGG enrichment analysis in D_vs_X. (D) Pathways-metabolite associated network.

### 3.3. Biomarker Identification based on AUC-ROC curve

A univariate analysis, ROC curve provided significant relation between D_vs_X with efficacy and generated AUC values (1 to 0.73). In this study, 247 DEMs AUC values were equal to 1 and 296 DEMs AUC values were closer to 1 (**Supplementary Table 4**). Metabolites AUC value equals to 1.0 represented a threshold for perfect separation between D_vs_X. Whereas, metabolites AUC value approaching to 1 provided more pronounced separation in D_vs_X and announced them as a potential biomarker. ROC curve analysis revealed that AUC of (-)-Riboflavin, Leukotriene D4, L-Glutamine were 0.973, FAD and Progesterone were 0.95, Fumaric acid, Glutathione disulfide, 15(S)-HETE, and Biliverdin were 0.89, and Glutathione was 0.73 which showed their role may be as a potential biomarker in ovarian physiology.

### 3.4. Integration of Metabolome and the Transcriptome

From transcriptome analysis for the same biological group, 195 DEGs were identified by using the parameters of fold Change≥1.5 and P-value<0.01, from which 108 were up-regulated and 87 were down-regulated **(unpublished data).** In order to identify metabolites that are potential regulators of gene expression in reproductive traits, we performed KEGG co-enrichment analysis between DEGs and DEMs. By adjusting the threshold of P-value<0.01 for DEGs and P-value<0.05 for DEMs, we obtained 48 KEGG pathways contributed by 85 DEMs and 37 DEGs. According to these results, determined the significantly enriched pathways including neuroactive ligand-receptor interaction, cAMP signaling pathway, apoptosis, insulin resistance, protein digestion and absorption, purine metabolism, glycine, serine and threonine metabolism, glutathione metabolism, ovarian steroidogenesis, drug metabolism - cytochrome P450, biosynthesis of amino acids, and metabolism of xenobiotics by cytochrome P450 (Supplementary Figure S2).

### 3.5. Correlation analysis, Cluster Analysis, and Correlation network between DEGs and DEMs

Log2 conversion data for the DEMs and DEGs used to calculate the correlation based on Pearson correlation method according to each difference grouping. The correlation between all genes and metabolites were selected that had a correlation coefficient (CC) > 0.8 and correlation p-value (CCP) <0.05 resulting into 34,936 corelated pairs. To obtain a systematic view of the variations in metabolites and their corresponding genes with CC and CCP, nine quadrant plot was generated (Figure 9A). In total, 1,048,575 gene-metabolite correlated pair were obtained through nine-quadrant plot. 73,337 and 66,335 gene and metabolite pairs were found in quadrant 3,7 and their expression trends were consistent which revealed that genes may be positively correlated with metabolites. 54,289, 45,253 gene-metabolite pair were found in quadrant 1, 9 and their expression trend were reversed which suggested that genes were negatively correlated with metabolites. 448804, 507,660 gene-metabolite pairs were noticed in quadrant 2, 8 with the upregulated and down regulated expression trend of metabolites and no change in genes expression. Interestingly, not a single gene-metabolite pair noticed in quadrant 4,5, and 6 (Figure 7A). Based on the differential metabolites and differential genes of each group, combined with the results of correlation analysis and heat map clustering analysis of the screened correlation revealed the expression level of each differentially expressed gene (195) and differentially expressed metabolite (542) in D_vs_X **(Figure 7B).** Then, based on K-mean classification we obtained two gene subcluster (no. of gene=108 and 87) and six metabolite subcluster (no. of metabolite in 1 & 2 subcluster=48; 3, 4, 5 and 6subcluster=120, 135, 67, and 124) **(Figure 7C) respectively.**

**Figure 7.**
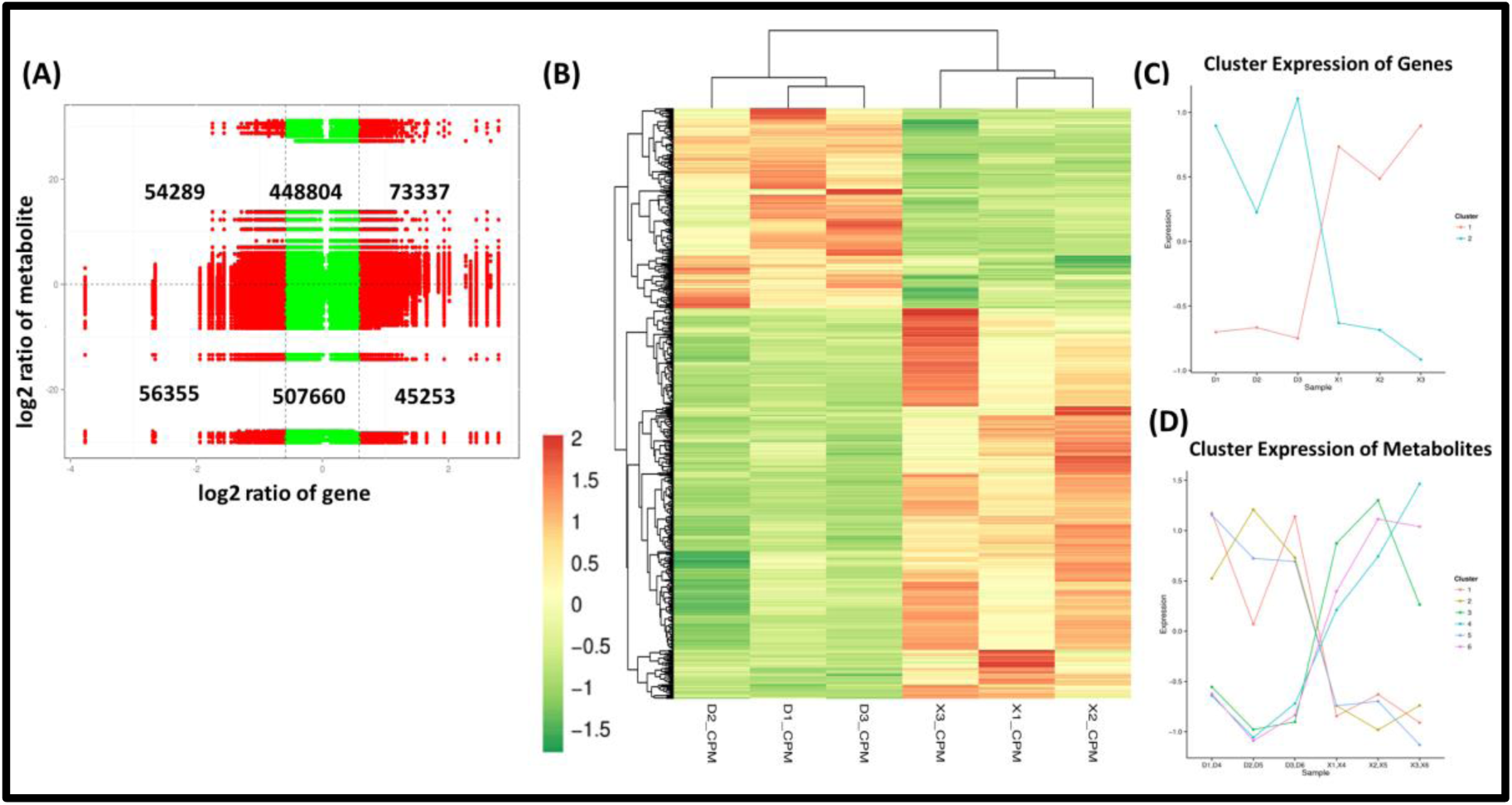
Gene-metabolite association based on Correlation coefficient (CC) and correlation p-value. (A) Scatter plot of 9-quadrant associate analyses of genes and metabolite from log2 FC. Number of points in each quadrant represented the number of gene and metabolites in each quadrant. parentheses. (B) Heat map of DEMs and DEGs expression pattern. (C) Differential gene and differential metabolite trend analysis.

Furthermore, linkage between metabolites and genes represented through the network diagram, and 20 differential genes and 47 differential metabolites screened by correlation, and selected for mapping along with their corresponding pathways **(Table 1 & Figure 8A-B)**. CORT exhibited positive correlation ((0.909) and C5AR1 exhibited negative correlation (−0.847) with Leukotriene D4 in the neuroactive ligand-receptor interaction pathway. Whereas, C3 positively correlated with Uridine 5’-diphosphate in neuroactive ligand-receptor interaction pathway. ATPP1A2 and PTCH1 were positively correlated with Succinic acid and L-(+)-Lactic acid contributed in cAMP signaling pathway. IFNGR1 was negatively correlated with L-(+)-Lactic acid involved in HIF-1 signaling pathway, and positively correlated with cortisone in cancer pathways. In folate biosynthesis, PGFS gene was positively correlated with the following metabolites; 2,5-diamino-6-hydroxy-4-(5-phosphoribosylamino)pyrimidine (0.822), Tetrahydrobiopterin (0.822), and 7,8-dihydroneopterin 3’-phosphate (0.859), whereas, negatively correlated with Guanosine triphosphate (−0.881). In the bile secretion pathway, we found positive correlation of PGFS with progesterone and ATP1A2 with Glutathione, Oxoglutaric acid, and Choline **(Table 1).** We observed PGFS was positively correlated with progesterone (0.821) in ovarian steroidogenesis pathway and negatively correlated with Leukotriene D4 (−0.893) in an arachidonic acid metabolism.

**Figure 8.**
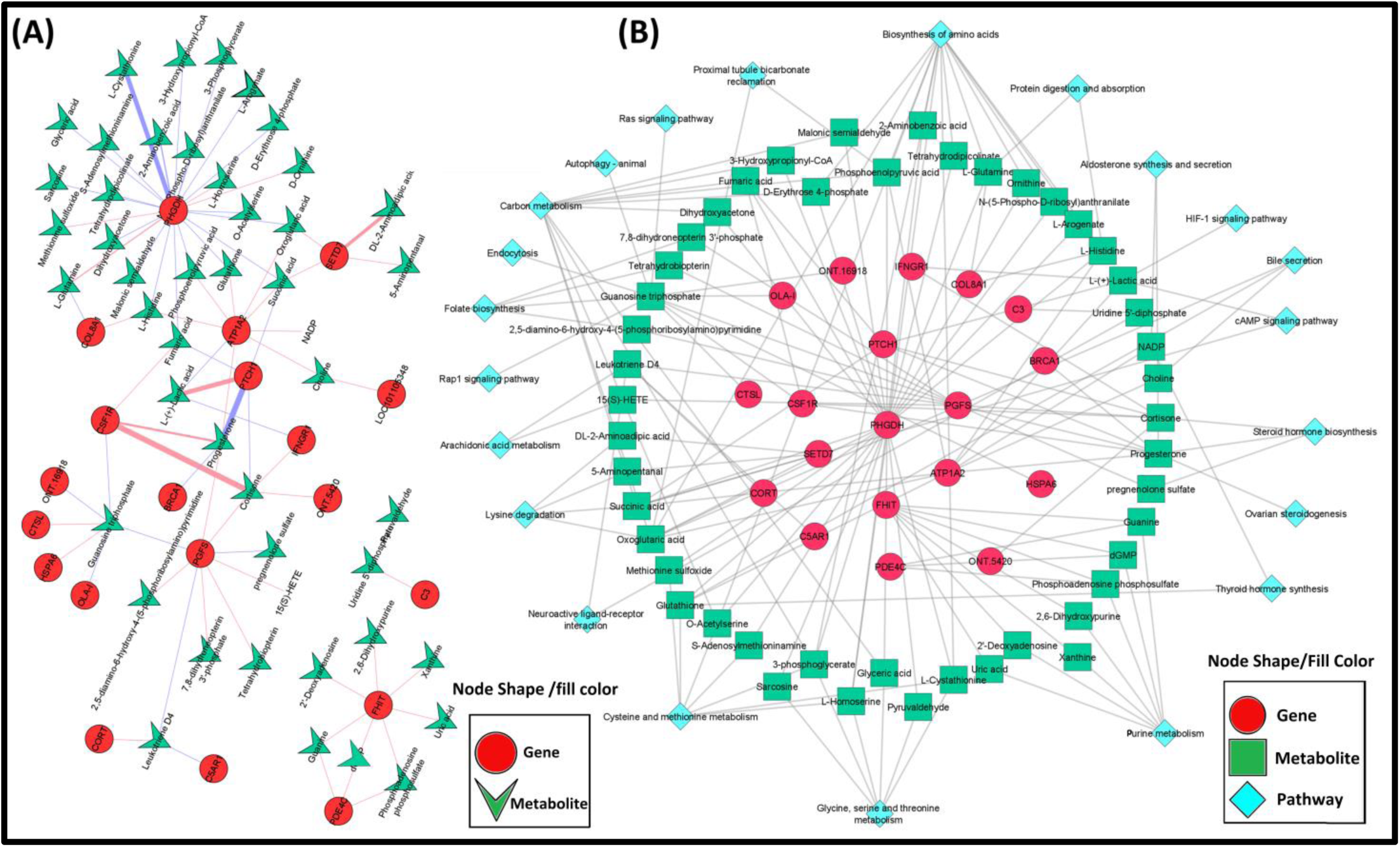
(A) Correlation based interaction network between genes and metabolites. The circle in the figure represents the gene, the triangle represents the metabolite. Note: The pink and blue line indicating positive correlation and negative correlation. Whereas, as larger the correlation coefficient, the wider the line, the darker the color. (B) Gene-Metabolite-Pathway interaction network.

**Table 1:**
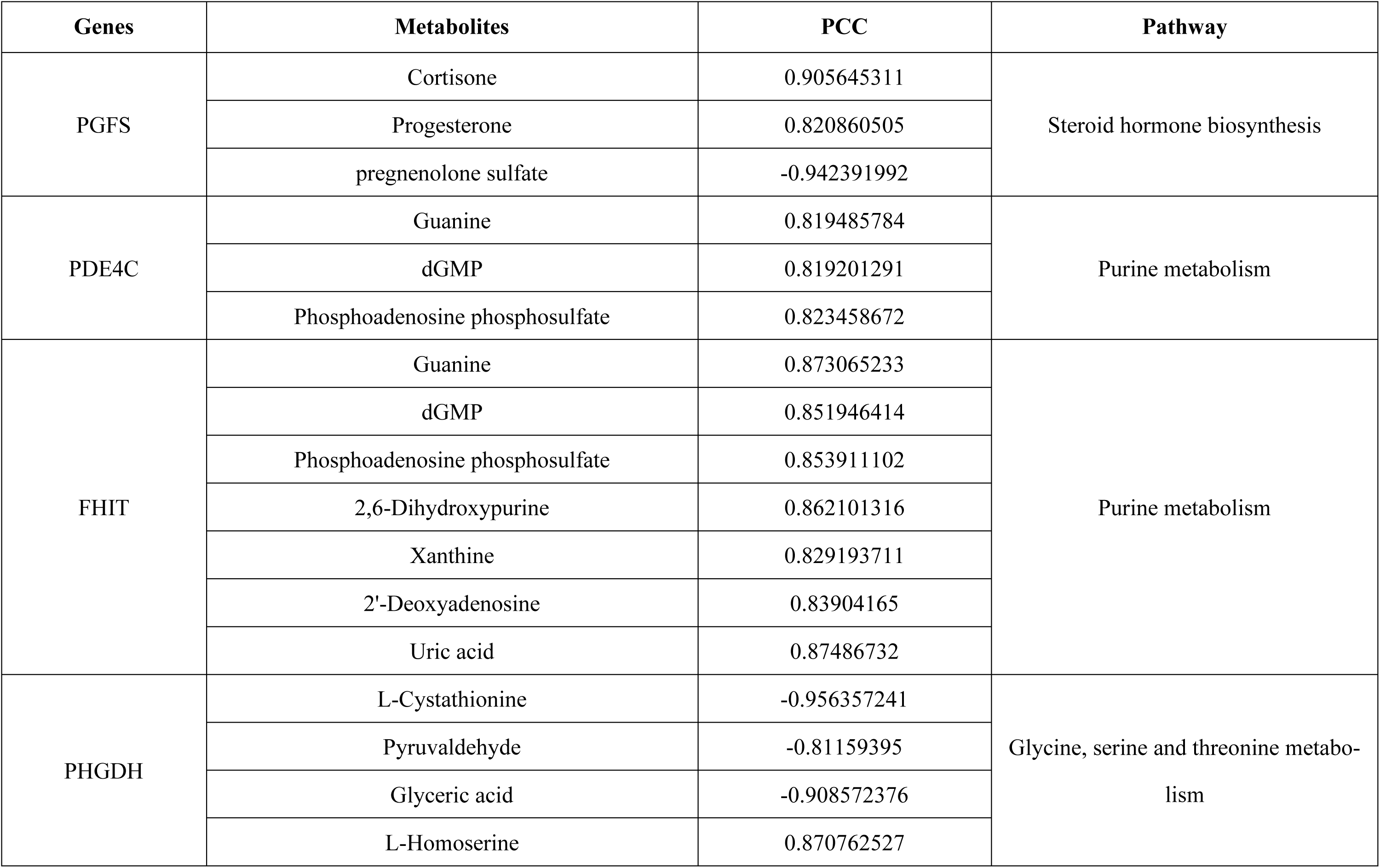
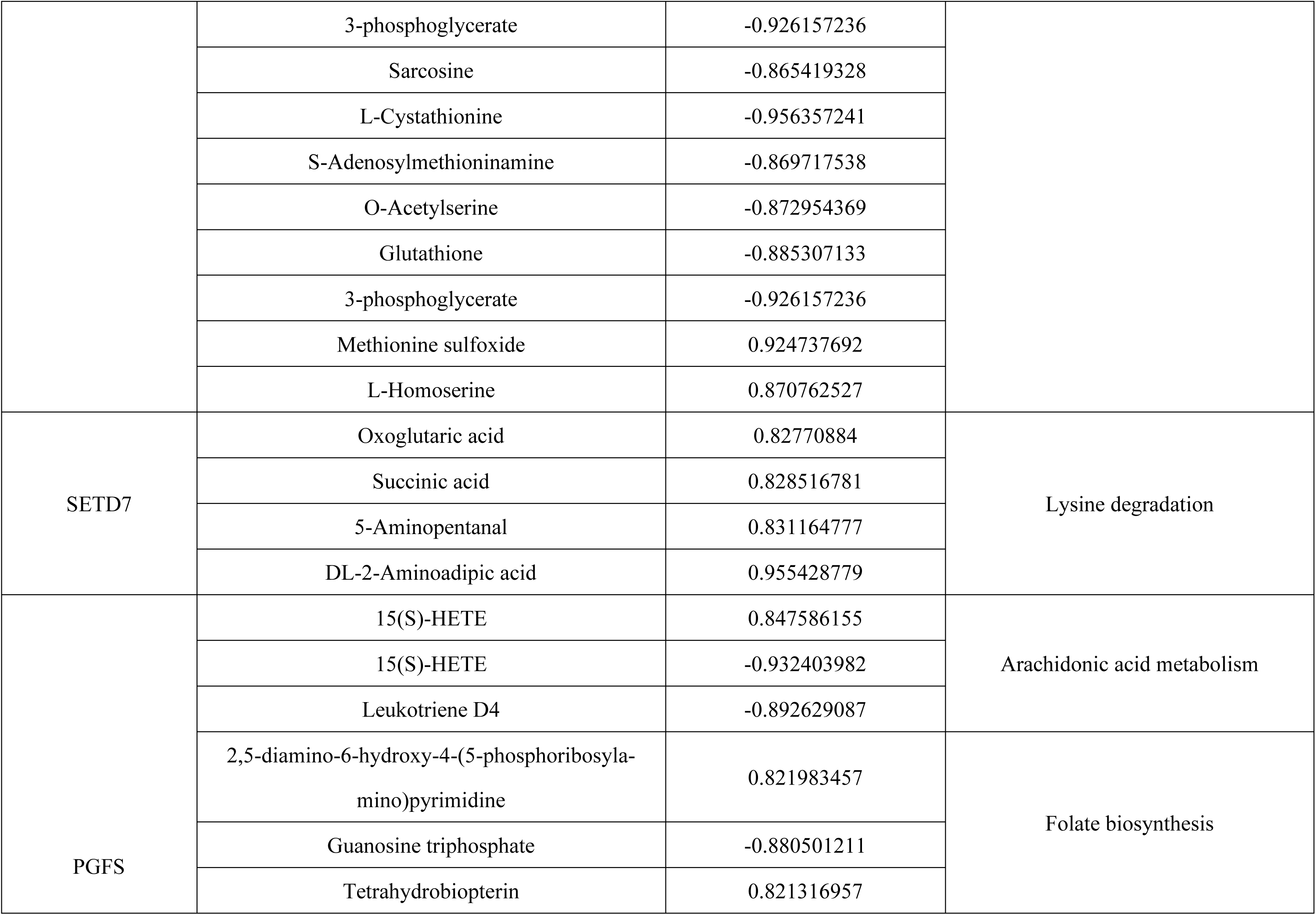
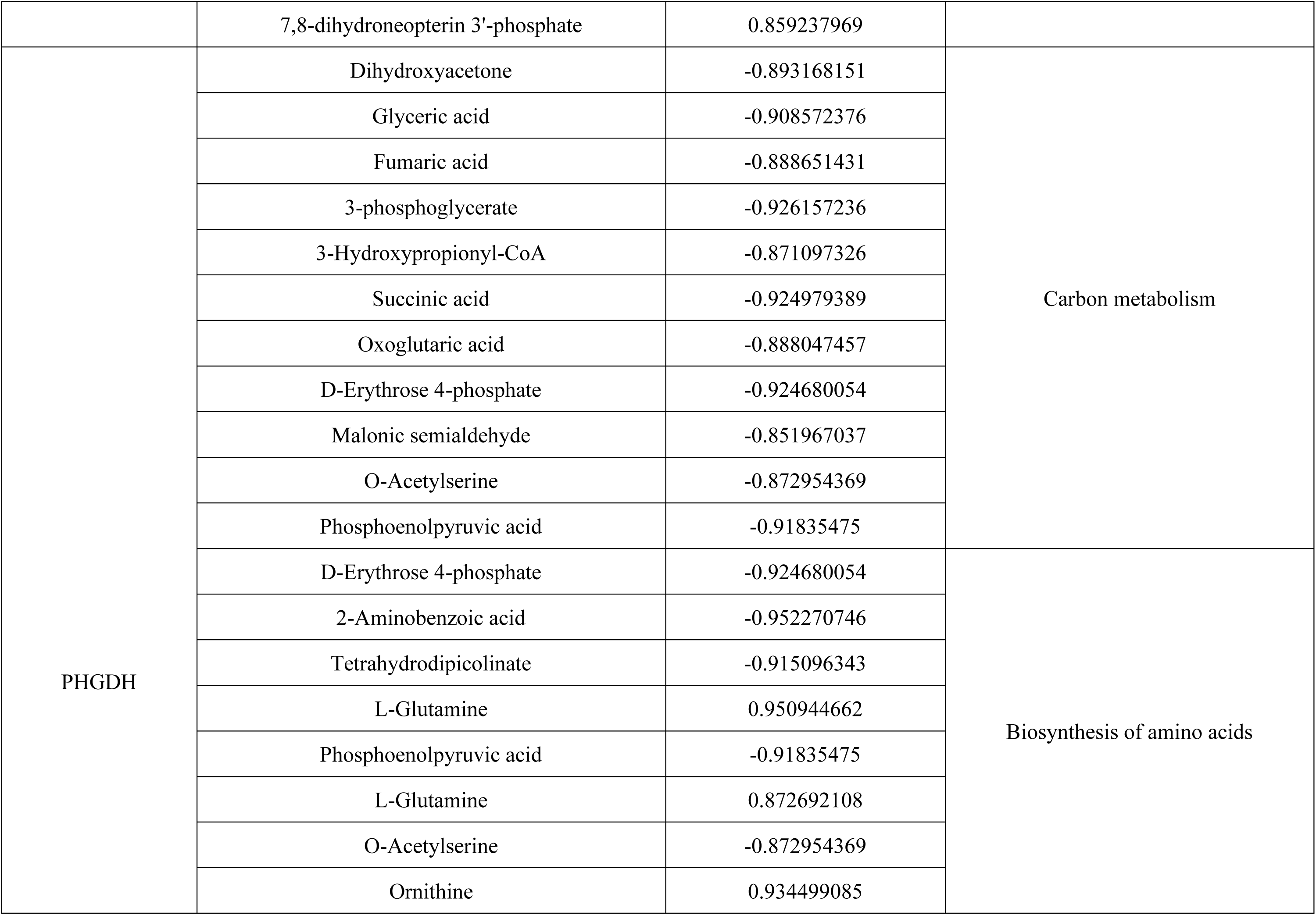
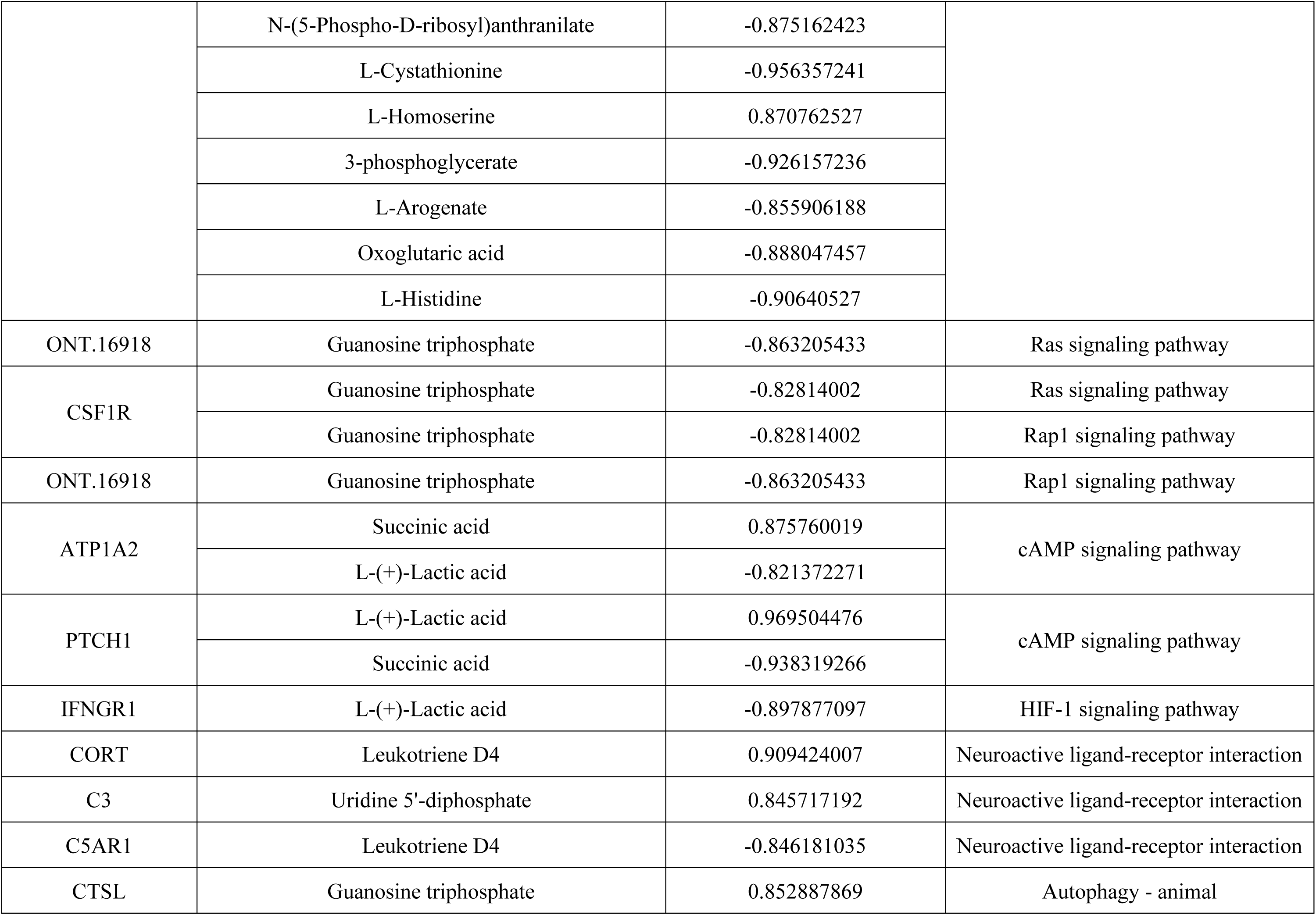
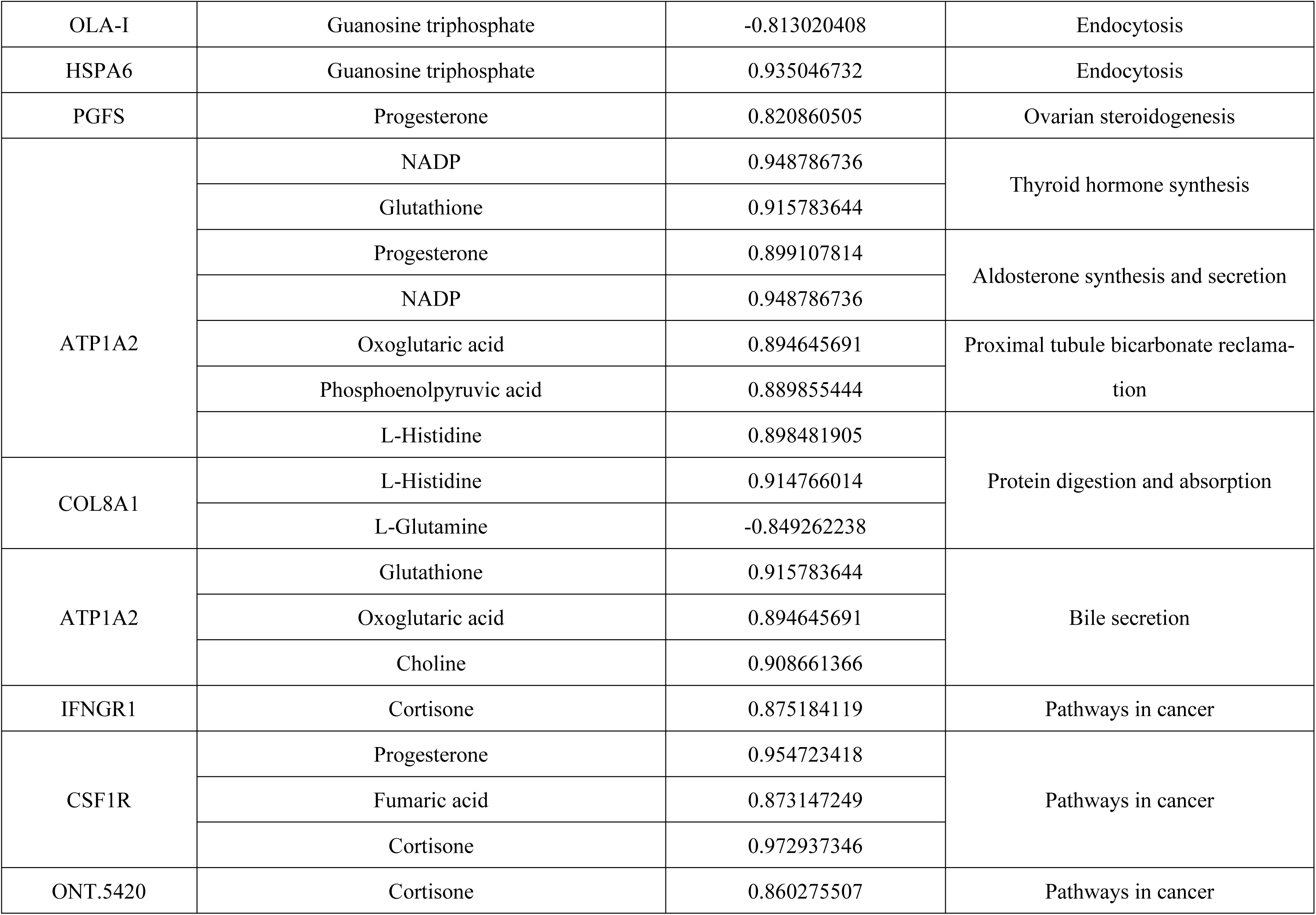
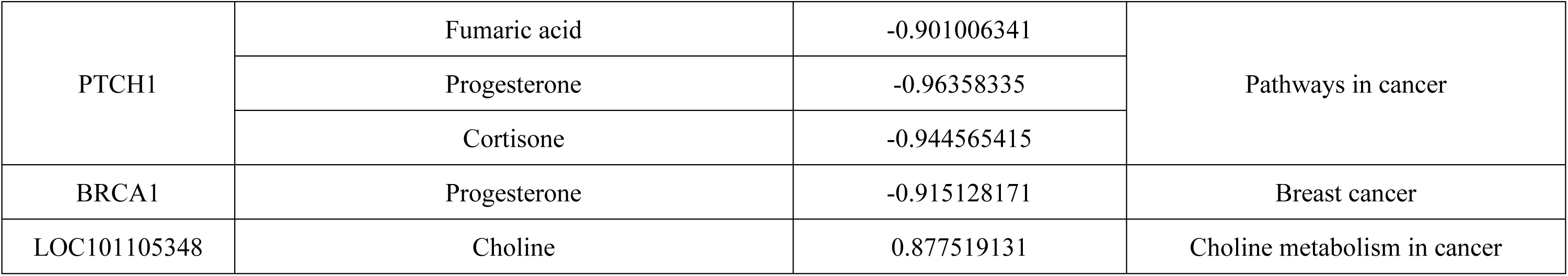
Gene-Metabolite association involved in significant pathways.

### 3.6. THE CANONICAL CORRELATION ANALYSIS

Canonical correlation analysis (CCA) established gene-metabolite association in neuroactive legend receptor interaction, protein digestion and absorption pathway, cancer pathway, purine metabolism, and cAMP signaling pathway. It can be visualized that PDE4C (gene8778), FHIT (gene22352) were correlated with Guanine (pos_6), dGMP (pos_293), Phosphoadenosine phosphosulfate (neg_444), 2,6-Dihydroxypurine (neg_27), Xanthine (neg_40), 2’-Deoxyadenosine (pos_204), and Uric acid (neg_44) in the pathway of purine metabolism (Figure 9A). ATP1A2 (gene1506) and PTCH1 (3095) were correlated with Succinic acid (neg_438) and L-(+)-Lactic acid (neg_5) in cAMP signaling pathway (Figure 9B). CSF1R (gene9993), IFNGR1(gene12728), ONT.5420, PTCH1 (gene3095) linked with Cortisone (neg_21), Progesterone (pos_728), and Fumaric acid (neg_473) in the biological process involved in regulating cancer (Figure 9C). COL8A1 (gene1868), ATP1A2 (gene1506) were correlated with L-Histidine (pos_489), and L-Glutamine (neg_108) in protein Digestion and absorption pathway (Figure 9D). C5AR1 (gene18421), CORT (gene15986), C3 (gene9228) were correlated with Leukotriene D4 (pos_130), Uridine 5’-diphosphate (pos_198) in neuroactive ligand receptor interaction pathway (Figure 9E).

**Figure 9.**
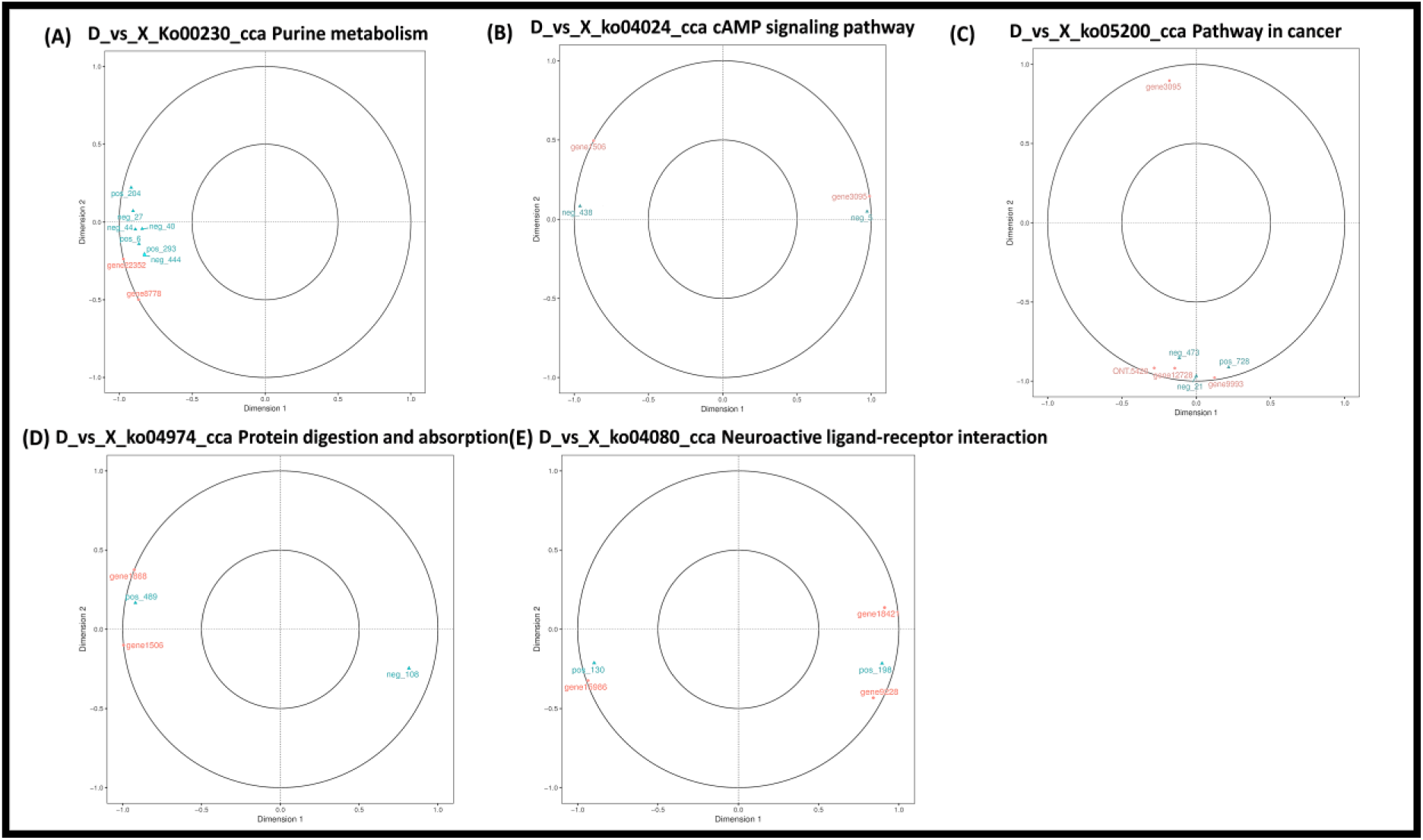
Typical correlation analysis plot and restrictive correspondence analysis (CCA) plot in D_vs_X. (A) Purine metabolism. (B) cAMP signaling pathway. (C) Protein digestion and absorption. (D) Neuroactive ligand-receptor interaction.

### 3.7. RESTRICTIVE CORRESPONDENCE ANALYSIS

Using a multivariate static correlation analysis with constrained correspondence, witnessed two restricted networks (**Figure 10A-C)**. C5AR1 (gene18421), CORT (gene15986), and C3 (gene 9228) were correlated with Leukotriene D4 (pos_130), Uridine 5’-diphosphate (pos_198) in neuroactive ligand-receptor interaction pathway, though, C3 (gene 9228) was positively correlated with Leukotriene D4 (pos-130), and negatively correlated with Uridine 5’-diphosphate (pos_198) **(Figure 10A).** Also observed that C5AR1 (gene18421) and CORT (gene15986) did not form a restricted network with any metabolites. On the other side, IFNGR1(gene12728) was correlated with Progesterone(pos_728), Fumaric acid (neg_473), Cortisone (neg_21) **(Figure 10C)**. Significant differences in the expression of genes and metabolites were noticed in D_VS_X as shown in Figure 12B-D, maybe they play crucial role in ovarian activity. In addition, (-)-Riboflavin, S-(Formylmethyl) glutathione, Fumaric acid, Leukotriene D4, and Progesterone AUC value 0.972, 0.916, 0.888, 0.972, 0.944 may indicate their potential use as biomarkers **(Figure 10 E-F-G-H-I-I)**.

**Figure 10.**
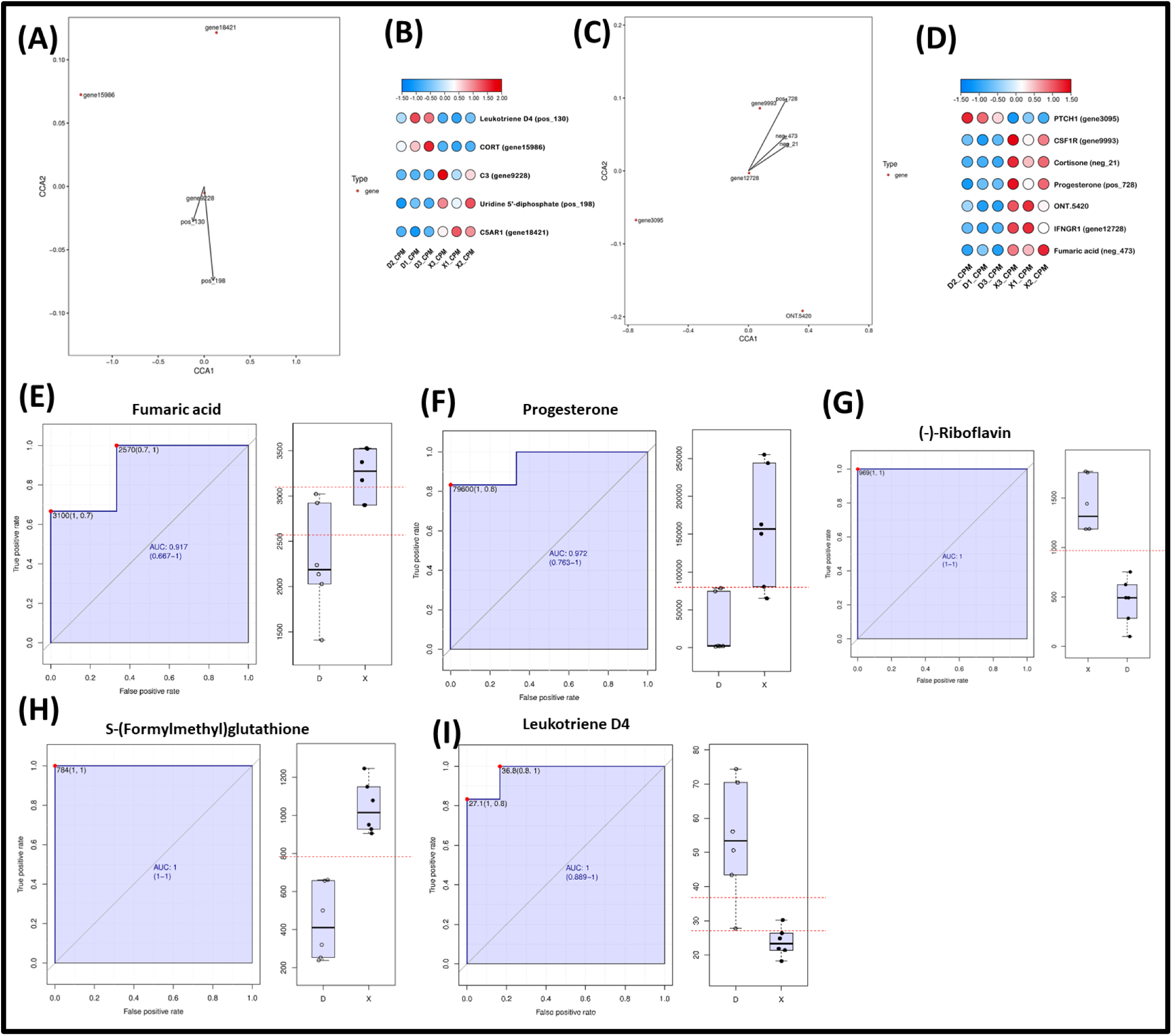
Restrictive correspondence analysis plot for gene and metabolite in respective pathway, their expression difference in D_vs_X and ROC-AUC curve analysis of predicted metabolite as a potential biomarker. (A-B) Neuroactive Ligand Receptor Interaction pathway. (C-D) Pathway in cancer. The dots in the figure indicate genes, and the arrows are metabolites. The angle between the gene and the origin line and the arrow reflects the correlation between the metabolite and the gene, acute angle being positively correlated and the obtuse angle being negatively correlated. (E-I) ROC-AUC curve analysis of fumaric acid, progesterone, and leukotriene D4 in D_vs_X was observed and determined as a potential biomarker.

## 4. Discussion

### 4.1. Significance of Differentially expressed metabolites in Sheep fecundity

Metabolites play compelling roles in biological process including reproduction, and their fluctuations impair fertility by affecting follicle health, growth, and development [51, 52]. Metabolic status impacts reproductive function at systemic level, modulating the hypothalamic GnRH neuronal network and/or the pituitary gonadotropin secretion through several hormones and neuropeptides, and at the ovarian level, acting through the regulation of follicle growth and steroidogenesis by means of the growth hormone-IGF-insulin system and local ovarian mediators [53]. Several studies have shown that biomolecules such as; lipids, amino acids, drugs and pollutants have a great impact on follicle growth and development [54], thus affecting fertility. In current study, we went through a comprehensive ovarian metabolic profiling in two sheep breeds with contrasting fecundity traits which provided us with 542 DEMs. Functional enrichment analysis and DEMs revealed their involvement in metabolic and signaling pathways in addition to determine significantly enriched pathways associated with reproduction such as; riboflavin metabolism and metabolism of xenobiotics by cytochrome P450, beta-Alanine metabolism, and secondary bile acid biosynthesis (Figure 7). Interestingly, several pathways resulting from DEGs and DEMs co-enrichment analysis were consistent with the KEGG enrichment analysis of DEMs such as; metabolism of xenobiotics by cytochrome P450, drug metabolism - cytochrome P450, neuroactive ligand-receptor interaction pathway, folate biosynthesis, and ovarian steroidogenesis as shown in Figure 6 & Supplementary Figure S1.

In current study Riboflavin metabolism was observed playing a significant role in traits associated with sheep reproduction and engaged following DEMs; (-)-Riboflavin (pos_4, neg_4), Guanosine triphosphate (pos_521) and FAD (neg_170) as shown in Figure 6D. Riboflavin, also known as vitamin B2, is an essential nutrient for livestock, including sheep, and plays a crucial role in energy metabolism, growth, reproduction, and antioxidant defense mechanisms. Riboflavin is a precursor of flavin mononucleotide (FMN) and flavin adenine dinucleotide (FAD), two important coenzymes involved in redox reactions, which are essential for ATP production, cellular respiration, and many metabolic processes [55]. Riboflavin metabolism has been shown to be closely associated with sheep fecundity, including ovulation rate, litter size, and reproductive performance[56].Previous studies also reinforce our findings, like pregnancy interruption due to deficiency of riboflavin was reported by [57, 58], their strong antioxidant roles mediated by free radicals was witnessed by [59]. In previous studies it has been observed the involvement of VB2 in ovarian development [60], bioenergetics pathways, redox homeostasis, chromatin remodeling [61, 62], deoxyribonucleic acid (DNA) repair mechanisms [63] and cell growth, and apoptosis [64, 65] which is in agreement to our findings of upregulated VB2 expression. ROC-AUC value determined that riboflavin (pos_4) may act as a potential biomarker for oxidative stress. Similarly, Guanosine triphosphate (GTP) and Flavin adenine dinucleotide (FAD) showed elevated expressions in current study which have been reported previously with implications in regulation of reproductive hormones (LH and FSH) in ovary cells [66–69] and reproduction associated energy metabolism [70] respectively. Moreover, Flavin adenine dinucleotide (FAD) is effective in various cellular redox reactions and the oxidative phosphorylation pathway providing sufficient oxygen supply for cellular activities and serves as a potential biomarker of metabolic activity in living cells [71]. Based on above evidences, we can infer that it may impair follicular growth and development and may provide a solid theoretical foundation to investigate the relationship between riboflavin metabolism and sheep fertility.

Xenobiotics are foreign chemicals either endogenous or exogenous which are modified through xenobiotic-metabolizing enzymes (XMEs) and involve in homeostasis and cellular integrity. Their metabolism has been observed significantly enriched in cytochrome P450 (CYP) pathway [72]. Previous studies demonstrated that the exposure of xenobiotics may destroy the primordial follicles which is responsible for premature ovarian failure and reduce fertility [49, 50]. In current study the following DEMs; S-(Formylmethyl) glutathione (pos_86, neg_134), Glutathione episulfonium ion (neg_86), and (1R)-Glutathionyl-(2R)-hydroxy-1,2-dihydronaphthalene (pos_411) were found involved in the xenobiotic metabolism *via* cytochrome p450 pathway (Figure 6D). All of these DEMs belong to glutathione (GSH) family, well known for their spectacular role as a free-radical scavenger, intervenient in xenobiotics metabolism, cell-cycle regulation, and a reservoir of cysteine [73]. GSH play key roles in cellular redox homeostasis [74]. A decreased glutathione disulfide level reflects oxidative stress due to damage in nucleic acid bases, lipids, and proteins [75] and depletion in glutathione cause premature ovarian aging [76], ovarian cyst [73] and infertility in female [77]. In addition, the protective action of follicle stimulating hormone on embryonic development is due to the synthesis of glutathione [78]. Another study reported the low level of glutathione reductase activity in reproductive aging [52]. Guo *et al.* determined the role of glutathione disulfide during ovulation in sheep ovaries, thereby participating in the metabolism of high-yielding ewes [79]. Eliasson *et al.* found the glutathione reductase cytosolic activities were significantly enhanced in the pregnant corpus luteum than the non-pregnant corpus luteum in pig [80]. In this study, mentioned metabolites of glutathione family were upregulated in small tail Han sheep and based on the above discussion we can infer that these metabolites have the potent role in ovary defense mechanism against exogenous and endogenous chemicals improving follicle fate and may act as biomarkers for treating anomalies related to ovarian function.

Notably, Beta-alanine is a non-essential amino acid that is found in various foods and is produced endogenously in animals via the degradation of carnosine [81]. In livestock, beta-alanine metabolism has been shown to play an important role in various physiological processes, including reproductive function in females. Specifically, beta-alanine has been shown to play a crucial role in the ovarian physiology of sheep, and it is believed that most beta-alanine metabolism occurs in the brain and muscles, and the final product of normal metabolism is acetic acid[82]. Meanwhile, beta-alanine can maintain the antioxidant capacity of the body [83], and the increase in beta-alanine could be an indicator of improved milk quality. Similarly, previous studies suggested that beta-alanine supplementation improved growth performance, meat quality, and antioxidant capacity, and also affected amino acid metabolism on broiler chickens [84], dairy cows [85]. One key role of beta-alanine in sheep ovarian physiology is in the regulation of follicular development, which is regulated by various hormones and growth factors. Beta-alanine has been shown to enhance the growth and development of ovarian follicles in sheep [86, 87], which can ultimately improve reproductive outcomes. The effects of alanine on reproductive traits in ruminants are still little discussed in the literature. However, it was demonstrated that, in an intrauterine environment with low nutrient and oxygen availability, sheep prioritized alanine and glutamine production for fetal organ growth and metabolism (a mechanism to sustain total energy needs [86]. Here, this study observed differences in alanine levels in two type of sheep breeds (Dolang sheep and Small Tail Han sheep), and found beta-alanine significantly enriched pathway in Small tail Han sheep, so, it can be suggested that the metabolites involved in beta-alanine metabolic pathway might be regulate small tail Han sheep fecundity through their nutritional status, and play crucial role particularly in the regulation of follicular development, luteal function, and reactive oxygen species (ROS) production. These findings suggest that beta-alanine supplementation may be a useful tool for improving reproductive outcomes in sheep and other livestock species. Further research is needed to fully elucidate the mechanisms underlying the effects of riboflavin metabolism, xenobiotics metabolism, and beta-alanine on reproductive function and to determine the optimal dietary interventions for improving reproductive performance in livestock.

### 4.2. Integrated Metabolomic and Transcriptomic Data Reveal Strong association between DEGs and DEMs In Regulating Reproduction Associated Traits

Metabolism is driven by specific enzymatic products of gene expression. In turn, gene expression requires the continual synthesis of certain metabolites and ATP. Thus, the two processes are coupled and must coordinate with one another as cells navigate altering conditions of existence [88]. Further, metabolites are present in metabolic pathways and their correlation with differentially expressed genes can better explain transcriptional regulatory mechanisms in metabolic pathways [89]. In this study, we mainly focused to investigate the association of DEGs and metabolites following new pathways involved in regulation of traits linked with sheep fertility. KEGG co-enrichment analysis provided us with **48** co-enriched pathways accompanied by **37** DEGs and **85** DEMs. These include variety of pathways related to reproduction like cAMP pathway, apoptosis, folate biosynthesis, oxytocin signaling pathway, ovarian steroidogenesis, endocytosis, glutathione metabolism, insulin resistance, drug metabolism-cytochrome p450, steroid hormone biosynthesis, Oocyte meiosis and neuroactive ligand receptor interaction pathway. Similarly, correlation-based gene-metabolite network analysis results into identification **20** DEGs and **49** DEMs involved in twenty-five biological pathways as shown in Figure 8A-B.

Existence of connectivity constraints in regulatory network determines the correlation between metabolites and gens [90]. Here, multivariate statistical analysis determined two constrained plots. In one plot, complement **component C3** (gene9228) exhibit positive correlation with **Leukotriene D4** (130) and **Uridine 5’-diphosphate** (pos_198) involved in **neuroactive ligand-receptor interaction pathway**. In second plot, **IFNGR1** (gene12728) interacted positively with progesterone (pos_728), fumaric acid (neg_473) and cortisone (neg_21) in cancer pathway along with significant implications in regulating various other metabolic pathways, including energy metabolism, drug metabolism, TCA cycle, Steroid metabolism (Figure 10 A-C). Several previous studies demonstrate the strong clues for these metabolites and pathways for being their involvement in regulation of fertility related traits at various levels. Like, Multiple transcriptome studies in animals such as goat [91], pig [92], poultry [93], zebrafish [94] have demonstrated the role of neuroactive ligand-receptor interaction pathway in reproductive activities. Zhang, T., et al reported the significant impact of neuroactive ligand-receptor interaction pathways and receptor protein tyrosine kinases on the regulation of folliculogenesis, oocyte maturation, and egg production [95]. Further, It has been witnessed previously that biologically active metabolites of arachidonic acid **“LTD4”** interact with chronic inflammatory conditions and affect reproductive efficiency in humans and in animals [96, 97]. **Uridine 5’**-diphosphate, known as pyrimidine ribonucleoside diphosphates, is an imperative cofactor in glycogenesis and perform crucial role in the function of Uridine 5’-diphospho glucuronosyltransferases (UDP-glucuronosyltransferases and improvement of ovarian immunity with positive effect on follicular growth and development [98]. Some studies reported complement system encoded by the C3 gene regulated at transcriptional level. This was produced in response to estrogen, and played an essential role in inflammation, phagocytosis, cellular immune responses [99–101], and important for implantation and maintaining pregnancy in mammals [101]. The results from current study are in agreement to these findings where the expression of C3 was downregulated, and the expression of Leukotriene D4 and Uridine 5’-diphosphate was upregulated in the D_*VS*_X (Figure 12B). Additionally, we found that Leukotriene D4 may act as a potential biomarker though AUC-ROC analysis (Figure.10 I). Therefore, C3 gene expression may induces effect on Leukotriene D4 and Uridine 5’-diphosphate in neuroactive ligand-receptor interaction pathway and play an important role in Small Tail Han sheep fecundity. Likewise, in current study, IFNGR1 gene has been observed associated with regulation of P4, fumaric acid, and cortisone metabolites in cancer pathway (Figure 10C) while **Progesterone** (P4) is an endogenous steroid sex hormone that supports reproductive function [87]. It is well known that progesterone plays a critical role in follicular development, maintenance of pregnancy in sheep [90] and cattle [91]. Furthermore, Progesterone is essential for the induction of GnRH synthesis before ovulation in postpartum ewes [94], associated with the increase of estrous fertility. Existing literature suggested that progesterone secretion declines with the ovarian ageing [102, 103], hence affecting reproductive traits. Here, we found the expression of P4, cortisone, fumaric acid, and IFNGR1 upregulated in Small Tail Han Sheep. Based on above discussion, their potent role in maintaining ovarian immunity, energy hemostasis and redox metabolism may be the reason for Small Tail Han Sheep higher prolificacy. The genetic alteration of IFNGR1 gene may lead to dysfunction of TCA, energy metabolism, steroid metabolism pathway and related enzymes, disrupt ovarian activity may induce ovary cancer, in turn, significantly impaired fertility. Additionally, progesterone and fumaric acid might act as a potential biomarker observed through AUC-ROC curve analysis (Figure 10E-F). Here, we conclude that IFNGR1association with progesterone, fumaric acid, and cortisone may essential for ovarian physiology and deviation may lead to metabolic disorder which impair fertility. Therefore, our result provides a solid theoretical foundation for investigation the association between genes and metabolites in sheep reproduction as well as to confirm their role as a potential biomarker in fertility disorders. All these mentioned pathways are important for their prominent role in reproduction. However, it is necessary to conduct more qualitative research to study profound effects of C3 on Leukotriene D4 and Uridine 5’-diphosphate, and IFNGR1 on Progesterone, cortisone, and fumaric acid effect on reproduction in sheep by gene editing technology, thereby investigating their roles as biomarkers in ovarian metabolic syndrome.

## 5. Conclusions

Based on the valuable findings of various researchers globally and results from current study, it could be concluded that the fertility of sheep is very complex and dynamic phenomenon. Reproductive traits have low heritability, discrete phenotypic expression and are expressed only in sexually mature ewes, leading to the selection of low intensity and long generational intervals. As metabolism is driven by specific enzymatic products of gene expression. In turn, gene expression requires the continual synthesis of certain metabolites and ATP. Thus, the two processes are coupled and must coordinate with one another as cells navigate changing conditions of existence. Hence, use of an alternative strategy of identification of key genes and their associated metabolites related to fecundity and their use as one of the selection criteria seems to be quite promising for improvement of fertility traits. This paper deals with an investigation at molecular level aimed to understand the role of key metabolites in fecundity of Small Tail Han Sheep. It aims to integrate metabolomics data with transcriptomic to show the association between gene and metabolite in specific pathway. The results revealed that **complement component *C3*** was significantly correlated with Leukotriene D4 and Uridine 5’-diphosphate in a Neuroactive ligand-receptor interaction pathway. ***IFNGR1*** was significantly correlated with Progesterone, Fumaric acid, and Cortisone in cancer pathway and regulating various metabolic pathways, including energy metabolism, drug metabolism, TCA cycle, and Steroid metabolism, thus influencing fecundity. The increased accuracy will ultimately pave the way for better economics of livestock farming particularly in small ruminants. Numbers of candidate genes affecting fertility have been found in many sheep breeds around the world. So constant genetic profiling of different breeds should be carried out in search for genes showing significant association with fecundity, fertility and prolificacy and their characterization should be done for better understanding of reproduction traits.

## 6. Patents

Not applicable

## Author Contributions

S.Y. performed the experiment, analyzed data and wrote the paper. X.Y.M. conceived and designed the study and wrote the paper. W.A.M, H.F., T.Y.L., and L.L.X. performed the experiment and interpreted the data. All the authors read and approved the final manuscript.

## Institutional Review Board Statement

All the procedures involving animals were approved by the animal care and use committee at the Institute of Animal Sciences, Chinese Academy of Agricultural Sciences (NO. IAS2019-82), where the study was conducted. All the experiments were performed in accordance with the relevant guidelines and regulations set by the Ministry of Agriculture of the People’s Republic of China.

## Informed Consent Statement

Not applicable

## Data Availability Statement

The materials and datasets used and analyzed during the present study are available from the corresponding author upon reasonable request.

