## Supplementary figures and images for "Integrated Transcriptomic and Metabolomic Profiling of Sheep Ovarian Tissues Confer their association in Fecundity associated Pathways"

### Supplemental Figure 1

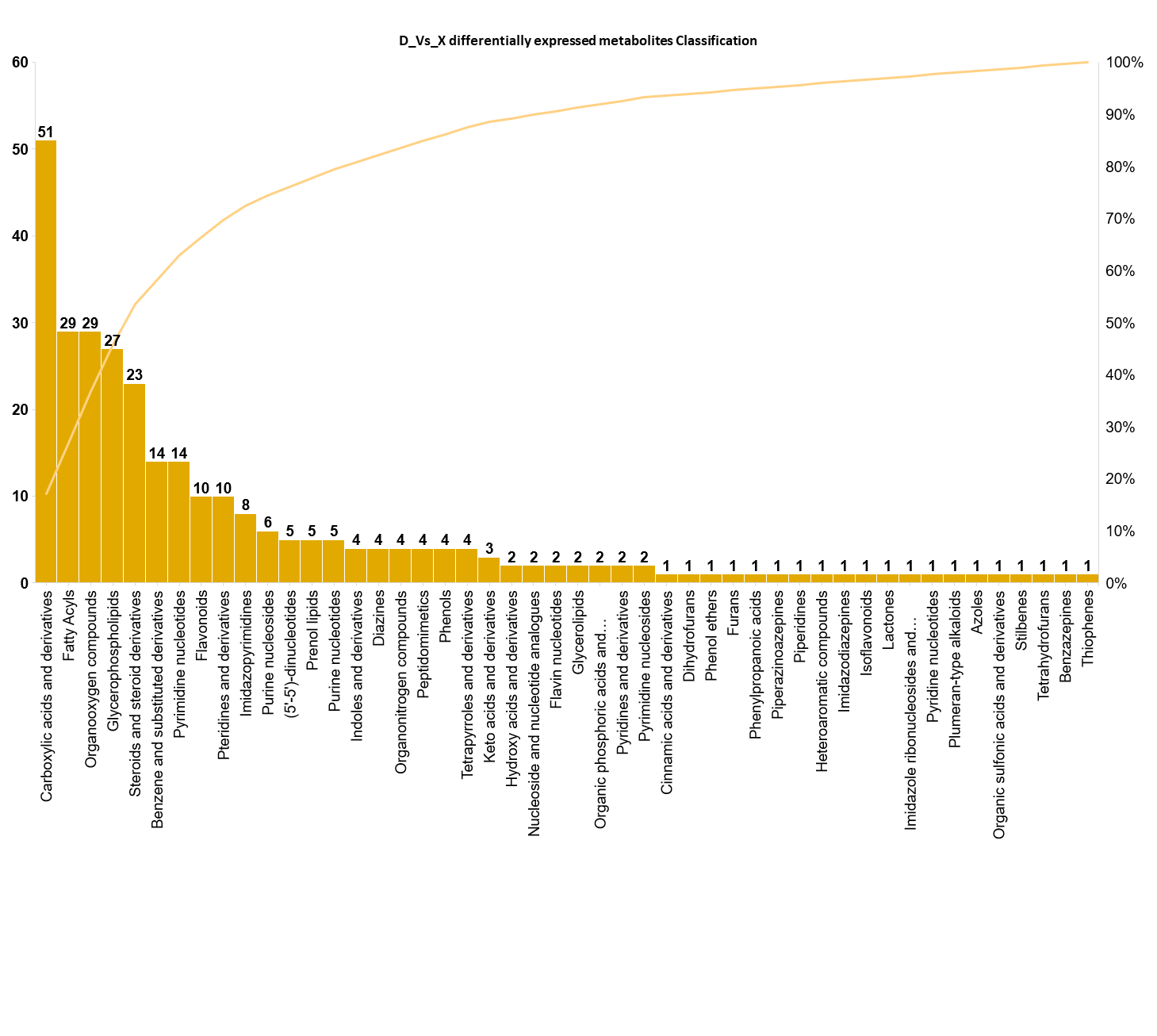

### Supplemental Figure 2

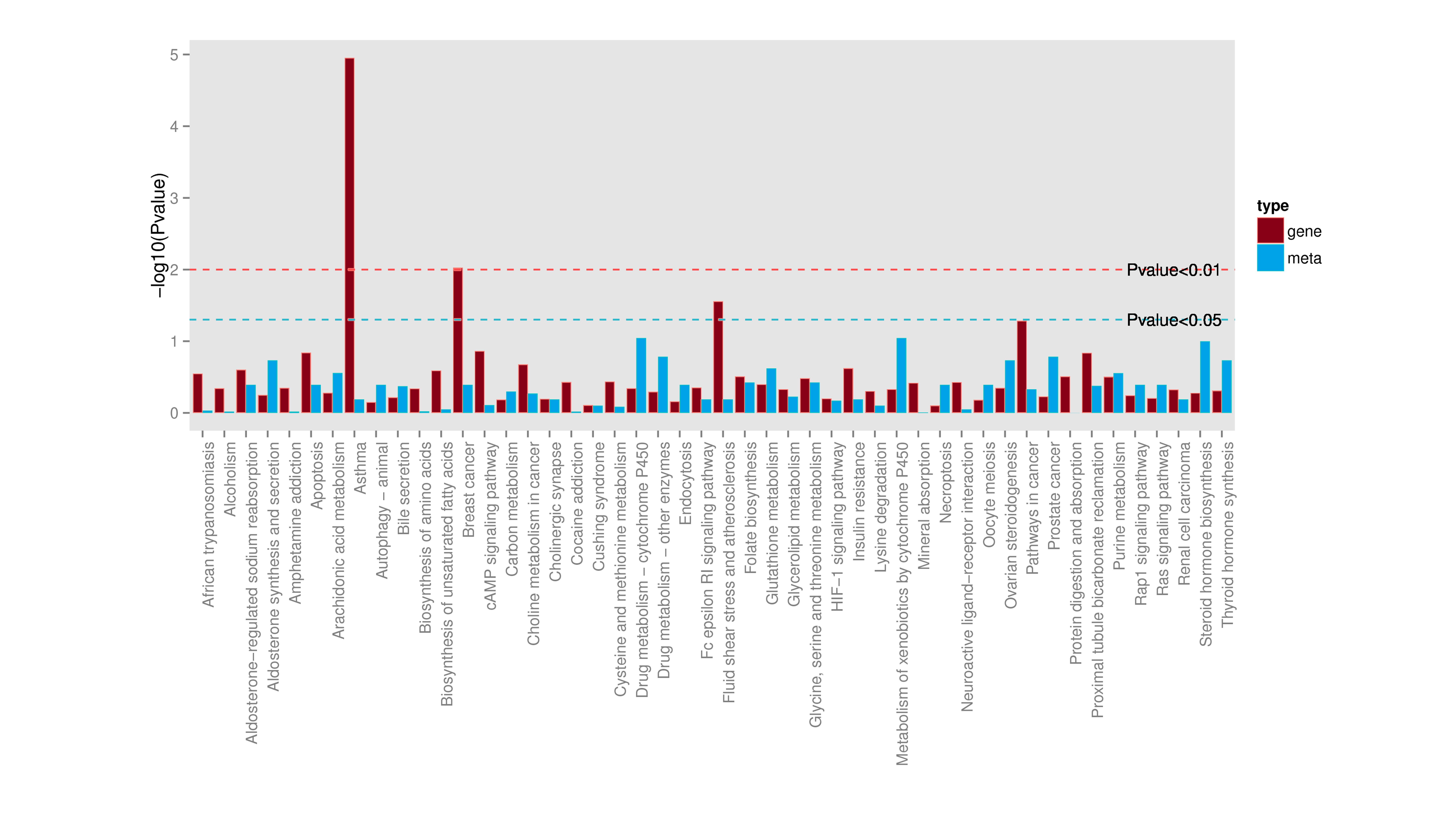
